# Genome-wide association analysis reveals new insights into the genetic architecture of defensive, agro-morphological and quality-related traits in cassava

**DOI:** 10.1101/2020.04.25.061440

**Authors:** Ismail Yusuf Rabbi, Siraj Ismail Kayondo, Guillaume Bauchet, Muyideen Yusuf, Cynthia Idhigu Aghogho, Kayode Ogunpaimo, Ruth Uwugiaren, Ikpan Andrew Smith, Prasad Peteti, Afolabi Agbona, Elizabeth Parkes, Ezenwaka Lydia, Marnin Wolfe, Jean-Luc Jannink, Chiedozie Egesi, Peter Kulakow

## Abstract

Cassava (*Manihot esculenta*) is one of the most important starchy root crops in the tropics due to its adaptation to marginal environments. Genetic progress in this clonally propagated crop can be accelerated through the discovery of markers and candidate genes that could be used in cassava breeding programs. We carried out a genome-wide association study (GWAS) using a panel of 5,310 clones developed at the International Institute of Tropical Agriculture - Nigeria. The population was genotyped at more than 100,000 SNP markers via genotyping-by-sequencing (GBS). Genomic regions underlying genetic variation for 14 traits classified broadly into four categories: biotic stress (cassava mosaic disease and cassava green mite severity); quality (dry matter content and carotenoid content) and plant agronomy (harvest index and plant type). We also included several agro-morphological traits related to leaves, stems and roots with high heritability. In total, 41 significant associations were uncovered. While some of the identified loci matched with those previously reported, we present additional association signals for the traits. We provide a catalogue of favourable alleles at the most significant SNP for each trait-locus combination and candidate genes occurring within the GWAS hits. These resources provide a foundation for the development of markers that could be used in cassava breeding programs and candidate genes for functional validation.

## 1.0 Introduction

Cassava (*Manihot esculenta* Crantz) is not only the most widely consumed starchy-root staple but also an emerging multi-purpose and industrial crop in Africa, Asia, and Latin America (Parmar et al. 2017). This clonally propagated species show remarkable adaptation to diverse agro-ecologies and can produce reasonable yield under marginal conditions of climate and soil (Jarvis et al. 2012). In addition, its flexible harvest window allows the crop to be left in the soil as a food reserve. These properties make cassava an ideal food security crop with an increasing trend in global production (Prudencio and Al-Hassan 1994; Burns et al. 2010).

In the last four decades, cassava breeding programs across Africa, Asia, and Latin America, have developed varieties that can withstand production constraints including biotic and abiotic stresses with improved yield and starch content (Kawano 2003; Okechukwu and Dixon 2008). While phenotype-based recurrent selection has made significant progress, the rate of genetic gain has been low due to several breeding complexities associated with the biology of the crop, including asynchronous flowering, low seed set per cross, a long cropping cycle of 12 to 24 months and low multiplication rate of planting materials (Ceballos et al. 2012). These challenges hinder the breeding program’s abilities to rapidly respond to changing human needs under volatile climatic and environmental conditions.

Modern breeding methods including marker-assisted selection (MAS) and genomic selection (GS) can be used to accelerate genetic improvement particularly by reducing generational interval and increasing selection intensity (Ferguson et al. 2012; Ceballos et al. 2015; García-Ruiz et al. 2016). However, integration of molecular markers as part of MAS in breeding pipelines requires an initial investment in discovery research to identify major-effect loci that serve as the targets of selection. With the rapid advances in next-generation sequencing (NGS) technologies, it is now feasible to generate genome-wide marker data in large populations. This, coupled with phenotype data makes it possible to identify and map locations of agriculturally important genes and quantitative trait loci (QTL) at the whole genome level (Varshney et al. 2014).

Significant investment has been made in the development of genomic resources for cassava including dense multi-parental linkage maps (International Cassava Genetic Map Consortium (ICGMC), 2015), annotated reference genomes (Prochnik et al. 2012; Zhang et al. 2018) and a haplotype map of common genetic variants from deep sequencing of hundreds of diverse clones (Ramu et al. 2017). Several genome-wide association studies (GWAS) have been conducted to describe the genetic architecture of resistance against cassava mosaic disease (Wolfe et al. 2016), reduced green mite infestation (Ezenwaka et al. 2018), cassava brown streak disease (Kayondo et al. 2018) and provitamin A and dry matter content (Esuma et al. 2016; Rabbi et al. 2017; Ikeogu et al. 2019).

Using a collection of 5,130 elite cassava clones derived from three cycles of recurrent selection in the International Institute of Tropical Agriculture (IITA) Cassava Breeding Program, we examined the genetic architecture of 14 continuous and categorical traits, including defense against biotic stresses, agro-morphological and quality-related traits (Table 1). Among the biotic stress-related traits, we considered cassava mosaic disease (CMD) and cassava green mite (CGM). Caused by different species of cassava mosaic geminiviruses, CMD is one of the most important biotic constraints to cassava production in Africa, India and Sri Lanka (Thottappilly et al. 2003; Alabi et al. 2015; CABI 2019) and has recently spread to the South Asian countries of Thailand and Vietnam (Uke et al. 2018). Infected plants can incur yield losses of up to 82% which translates to more than 30 million tonnes of fresh cassava roots loss annually (Owor et al. 2004; Legg et al. 2006). Infestation by CGM (*Mononychellus tanajoa*) during the dry season causes chlorosis and restricted growth resulting in a significant negative impact on root yield. The main type of resistance to CGM is attributed to apical leaf pubescence but may also include other mechanisms (Shukla 1976; Byrne et al. 1982; Ezenwaka et al. 2018).

**Table 1.** Name, brief description, and classification of the 14 traits assessed in the present study

| Name of Trait | Description | Class |
| --- | --- | --- |
| Cassava mosaic disease (CMD) severity | The visual rating of symptoms caused by cassava mosaic virus | Biotic stress |
| Cassava green mite (CGM) severity | The visual rating of damage caused by cassava green mite | Biotic stress |
| Apical pubescence | Pubescence of young apical leaves | Morphological |
| Leaf shape | The shape of central leaf taken from a mid-height position. | Morphological |
| Apical leaf colour | Colour of unexpanded apical leaves | Morphological |
| Mature leaf greenness | Colour of first fully expanded leaf, an indicator of leaf chlorophyll content | Morphological |
| Petiole colour | Pigmentation of leaf petioles | Morphological |
| Harvest index | Harvest index | Agronomic traits |
| Plant type | Plant architecture on a 1-5 scale | Morphological |
| Outer stem colour | Stem colour nine months after planting | Morphological |
| Total carotenoid (colour chart) | Level of yellowness in cassava storage root pulp (parenchyma) due to variation in carotenoid content | Quality traits |
| Dry matter content | Percentage of dry matter content of storage roots | Quality traits |
| Storage root periderm colour | Colour of the outer surface of storage root periderm (outer skin) | Morphological |
| Storage root cortex colour | Colour of the outer surface of storage root cortex (inner skin) | Morphological |

For quality traits, we considered total carotenoid content and dry matter content variation. Biofortification breeding to increase provitamin A carotenoids in storage roots is an important goal in cassava improvement programs around the world (Saltzman et al. 2013). Although the crop’s gene pool contains accessions that are naturally rich in provitamin A, they make up a small proportion of cultivated varieties especially in Africa (Welsch et al. 2010). We used the colour-chart based assessment of total carotenoid content variation. In cassava, the intensity of root yellowness is strongly correlated with total carotenoid content (Sánchez et al. 2014). Additionally, provitamin A is the predominant carotenoid component in cassava (Ceballos et al. 2017). Dry matter content is a key yield component that determines varietal acceptance by farmers and processors (Bechoff et al. 2018). Cassava germplasm contains considerable variation in percentage dry matter content of the fresh roots ranging from as low as 10% to 45% (Kawano et al. 1987).

Among agronomic traits, harvest index and plant type were included in the present study. Harvest index reflects the partitioning of resources between the storage roots and above-ground biomass (Sinclair 1998). The desirable harvest index values for the crop range from 0.5 to 0.7 (Kawano et al. 1978; Kawano et al. 1998). Plant type in cassava can be characterized by four general descriptive shapes, namely cylindrical, umbrella, compact, and open. Plants with cylindrical shapes do not form branches and are most preferred for mechanized farming. Umbrella plant types generally branch at a high level (above 1 m) and have fewer levels of branching whereas compact and open types are characterized by low first branching height and multiple branching levels but differ in the angle of branches and erectness of the stems (Fukuda et al. 2010).

We also assessed the genetic architecture of several morphological traits related to leaves (petiole colour, apical leaf colour, mature leaf greenness, leaf shape) and storage roots (periderm and cortex colour). Colour of cassava leaf petioles as well as apical leaves ranges from light green to purple due to anthocyanin pigmentation. Anthocyanins play various roles in plants including protection against ultraviolet light, overcoming different abiotic and biotic stresses and other physiological processes such as leaf senescence (Gould et al. 2008). Mature leaf greenness is related to chlorophyll content, an indicator of a plant’s photosynthetic capacity (Palta 1990). Additionally, cassava germplasm exhibits diverse leaf shapes ranging from ovoid lobes to linear forms (Fukuda et al. 2010). This variation is useful as a morphological descriptor and could also play other functional role related to light capture (Takenaka 1994). Cassava root periderm colour varies from cream through light brown to dark brown while that of the cortex includes cream, pink and purple. The few but predominant farmer-preferred varieties in Africa often have a pink or purple cortex and dark brown periderm although there is no proven genetic correlation between these traits and the culinary properties in cassava. On the other hand, industrial processing into starch and flour production generally utilize whole roots after mechanical periderm removal. These industries prefer varieties with white/cream periderm in order to ensure bright-coloured products. Finally, we also assessed the genetic architecture of stem colour, another morphological descriptor used for variety identification and varies from orange to dark-brown (Fukuda et al. 2010).

Here, we provide a catalogue of genomic loci associated with the 14 traits, a list of favourable alleles for each trait-locus combination and candidate genes located within the identified loci. Understanding the genetic architecture of variation in the studied traits is an important step towards the development of molecular tools to accelerate the transfer of favourable alleles into farmer-preferred varieties.

## Material and Methods

### Field experiments and phenotyping

The breeding population composed of 5,130 elite IITA cassava breeding genotypes were phenotyped for the 14 traits across four contrasting locations in Nigeria; Ubiaja (6°40’ N, 6°20’ E), Ibadan (7°24’ N, 3°54’ E), Mokwa (9°21’ N 5°00’ E) and Ikenne (6°52’ N 3°42’E) from 2013 to 2016. Description of the traits, their ontologies and measurement methods are available at https://cassavabase.org/search/traits. This population consisted of 717 elite lines from the genetic gain (GG) population, 2,322 full-sib progenies derived from 88 elite GG progenitors (Cycle1 – C1), and 2,091 full-sib progenies derived from top 89 cycle one progenitors (Cycle2 – C2) (Supplementary Table S1). The mean family size for C1 clones was 15, ranging from 1 to 77 clones while that of C2 was 7.6 ranging from 1 to 20 clones. The GG population is a collection of the “Tropical *Manihot* Selection (TMS)” cultivars developed in the past five decades by IITA (Dixon and Ssemakula 2008; Rabbi et al. 2017). This panel represents an extensive and diverse pedigree comprising of crosses between West African landraces, Latin American elite and wild germplasm. The raw phenotypic data is openly accessible on CassavaBase repository (ftp://ftp.cassavabase.org/manuscripts/PlantMolBiol_Rabbi_et_al_2020/)

The GG population was evaluated in clonal evaluation trials (CET) while the C1 and C2 were evaluated in CET, preliminary yield trials (PYT), and advanced yield trials (AYT). The CET plots were composed of a single row with 5 plants per plot, in an augmented block design, with 2 checks as a control. PYT plots consisted of two rows with 10 plants per plot, in a randomized complete block design with 2 reps and 5 checks as a control. AYT plots contained four rows with 20 plants per plot, in a randomized complete block design with 3 reps and 5 checks as a control. Planting was performed from June to July (during the rainy season) and harvested around the same time the following year. Spacings between rows and plants were 1 and 0.8 m in all trials, respectively, except in CETs where we used 0.5 m within rows. All field trial management was performed, whenever necessary, in accordance with the technical recommendations and standard agricultural practices for cassava (Fukuda et al. 2010; Abass et al. 2014; Atser et al. 2017).

### Genotyping

Genomic DNA was extracted from freeze-dried leaf samples following a modified Dellaporta CTAB method (Dellaporta et al. 1983). DNA quality and quantity were assessed using a Nanodrop 1000 spectrophotometer at 260 nm absorbance. Genome-wide single nucleotide polymorphism (SNP) data was generated using the genotyping-by-sequencing (GBS) approach described by Elshire et al. (2011). Reduction in genome complexity for GBS was achieved through restriction digestion using ApeKI enzyme (Hamblin and Rabbi 2014). Sequencing reads were aligned to the cassava V6 reference genome (Prochnik et al. 2012) followed by SNP calling using TASSEL GBS pipeline V4 (Glaubitz et al. 2014). SNP calls with less than 5 reads were masked before imputation using Beagle V4.1 (Browning 2016). A total of 202,789 biallelic SNP markers with an estimated allelic r-squared value (AR2) of more than 0.3 were retained for subsequent analyses after imputation.

## Statistical analyses

### Phenotype data analyses

Our interest in this study was to identify the genetic architecture of selected traits rather than location- or year-specific QTLs. We collapsed plot observations for each genotype to a single best linear unbiased prediction (BLUP) using the following mixed linear model (MLM) with the lme4 (Bates et al. 2015) package in R (R Development Core Team 2016): *y_ij_* = *μ* + *g_i_* + *β_j_* + *r*_*j*(*l*)_ + *ε_ijl_* where *y_ij_* represents vector of phenotype data, *μ* is the grand mean, *g_i_* is the random effect of genotype i with 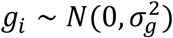, *β_j_* are the fixed effects of year – location combination j, *r*_*j*(*l*)_ is a random effect of replication nested within location-year combination assumed to be distributed 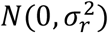; and *ε_ijl_* is the residual with 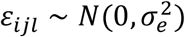.

Pairwise correlation between traits was determined from BLUPs using the R function “*cor*” in the “*stats*” package (R Development Core Team 2016) and visualization of the correlation matrices was done using the ‘*corrplot*’ R package (Wei and Simko 2017). Due to the unbalanced nature of trials, we calculated the broad-sense heritability estimates on a plot-mean basis using the formula 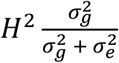. Additive genomic heritability for each trait was also estimated using a linear mixed model as implemented in GCTA (Yang et al. 2011; Speed et al. 2012).

### Population structure and genetic relatedness assessment

Population structure and cryptic genetic relatedness are known to confound GWAS analysis and lead to spurious associations (Chen et al. 2016). To assess the extent of population structure, we performed a principal component analysis (PCA) using PLINK-v1.9 (Purcell et al. 2007; Rentería et al. 2013). The pairwise Kinship matrix was visualized using the heatmap function in R.

### Genetic architecture of studied traits

For each trait, single-locus GWAS analysis was carried out using the mixed linear model (MLM) approach implemented in Genome-Wide Complex Trait Analysis (GCTA) software (Yang et al. 2011). Because the inclusion of the candidate marker in kinship calculation when controlling for cryptic relatedness can lead to a loss in power (Yang et al. 2011; Listgarten et al. 2012), we also used the mixed linear model which excluded candidate markers via a leave-one-chromosome-out analysis as implemented in GCTA (Yang et al. 2011; Yang et al. 2014). The first approach is referred to as MLMi (“i” for candidate marker included) and the second approach is referred to as MLMe (“e” for candidate marker excluded). Visualisation of MLMi and MLMe results in the form of Manhattan and quantile-quantile plots were implemented in the ‘*CMplot*’ R package (LiLin-Yin 2019).

### SNP Marker and favourable allelic prediction

To assess the genetic architecture of the studied traits, we fit a linear regression model using peak SNPs at the identified loci as independent variables against the traits’ BLUPS as the response variables. The relative allele substitution effects at each marker were visualized using boxplots. Here, a locus was defined as a uniquely identifiable genomic region whose SNPs passed genome-wide Bonferroni significance threshold (α=0.05/101,521 = 4.93 × 10^-7^).

### Candidate gene identification

Candidate loci were explored using a combination of GWAS p-values, local linkage disequilibrium (measured as <u>r^2^</u>), and gene annotation using the gff3 file of the cassava genome available on phytozome v.12.1 (https://phytozome.jgi.doe.gov/pz/portal.html) (Goodstein et al. 2011; Batra et al. 2014). This information set was summarized for each candidate loci using Locus zoom (Pruim et al. 2010). To obtain a regional zoom plot for each candidate locus, we built a local SQLite database including the 101,521 biallelic SNP marker matrix, associated GWAS p-values for each trait analyzed and the cassava gene-annotation following instructions available at https://genome.sph.umich.edu/wiki/LocusZoom_Standalone. Gene codes were shortened to ease visualization and whenever available, *Arabidopsis thaliana* homologues were noted. Recombination information was provided using the same approach as described in Wolfe et al. (2016). A standard interval of 100 Kb (50 kb upstream, 50 kb downstream) was explored for each candidate locus and adjusted according to the extent of local linkage disequilibrium with the candidate SNP (*r*^2^>0.8).

## Results

### Variation and relationships among traits

The frequency distributions of the BLUPs per trait within the panel are presented in Supplementary Figure S1 and the raw BLUPs values are also presented in Supplementary Table S1. An analysis of the phenotypic classes of the panel showed that almost all measured traits followed a normal distribution. A couple of traits were slightly skewed towards the tails. The heritability and variance components estimates associated with the studied traits are presented in Table 2. All measured traits exhibited larger than 2 fold differences between the maximum and minimum values with a mean coefficient of variation (CV) of 35%. Apical pubescence was the only trait with a significantly larger CV of 113%. These large differences between the maximum and minimum values are an indication of broad genetic variability within the mapping panel. To estimate the influence of additive and non-additive genetic effects on the observed phenotypic values, we estimated broad-sense and genomic-heritabilities. Both the broad-sense and SNP-based heritability estimates were comparable ranging from low to high: 15 to 78% and 17 to 72%, respectively. Some traits like total carotenoids, dry matter, petiole colour, apical leaf colour, and stem colour had higher SNP-heritability estimates (> 0.5) relative to other traits like harvest index, plant type, and resistance to cassava green mite with the lower heritabilities (< 0.4). SNP-based heritability is useful in approximating the proportion of phenotypic variance attributable to the additive genetic variation (Yang et al. 2017). The moderate to high levels of SNP-based heritabilities found for traits under active selection coupled with sufficient variability in the population indicates good potential for genetic improvement of these traits through recurrent selection.

**Table 2.** Broad-sense and SNP heritability estimates, variance components and coefficients of variation of 14 traits in cassava GWAS panel

| Trait | SNP-h<sup>2</sup> | H<sup>2</sup> | $\sigma_g$ | $\sigma_{g \times e}$ | $\sigma_e$ | CV |
| --- | --- | --- | --- | --- | --- | --- |
| CMD severity | 0.434 | 0.776 | 0.783 | 0.086 | 0.140 | 63 |
| CGM severity | 0.165 | 0.149 | 0.074 | 0.177 | 0.244 | 19 |
| Apical pubescence | 0.502 | 0.531 | 0.175 | 0.083 | 0.071 | 113 |
| Leaf shape | 0.499 | 0.510 | 0.679 | 0.475 | 0.178 | 22 |
| Apical leaf colour | 0.496 | 0.567 | 1.610 | 0.629 | 0.601 | 23 |
| Mature leaf greenness | 0.568 | 0.531 | 0.427 | 0.186 | 0.191 | 18 |
| Petiole colour | 0.716 | 0.754 | 3.322 | 0.602 | 0.485 | 33 |
| Harvest index | 0.308 | 0.538 | 0.010 | 0.002 | 0.007 | 28 |
| Plant type | 0.384 | 0.369 | 0.376 | 0.180 | 0.465 | 38 |
| Outer stem colour | 0.516 | 0.388 | 0.907 | 0.191 | 1.241 | 22 |
| Total carotenoids content (colour chart) | 0.675 | 0.726 | 0.401 | 0.066 | 0.085 | 49 |
| Dry matter content | 0.565 | 0.521 | 14.776 | 3.385 | 10.184 | 15 |
| Root periderm colour | 0.548 | 0.610 | 0.190 | 0.035 | 0.086 | 19 |
| Root cortex colour | 0.518 | 0.415 | 0.070 | 0.036 | 0.062 | 30 |

Where: H<sup>2</sup> is the Broad-sense heritability, $\sigma_g$ is the clonal genotypic variance, $\sigma_{g \times e}$ is the variance due to genotype by environment (G×E), and $\sigma_e$ being the residual variance.

The relative magnitude of phenotypic correlation pairs ranged from 0.27 between total carotenoid variation by colour chart (TC-chart) and root cortex colour to −0.65 for mature leaf greenness and petiole colour (Figure 1). We detected highly significant negative phenotypic correlations between dry matter and total carotenoid contents (−0.31, P < 0.001); mean CMD severity and harvest index (r = −0.14, P < 0.001); and apical pubescence and CGM severity (r = −0.27, P < 0.001). There were highly significant positive correlations between total carotenoids and periderm colour (r = 0.27, P<0.001), leaf greenness (r = 0.17, P < 0.001), and stem colour (r = 0.12, P < 0.001).

**Figure 1.**
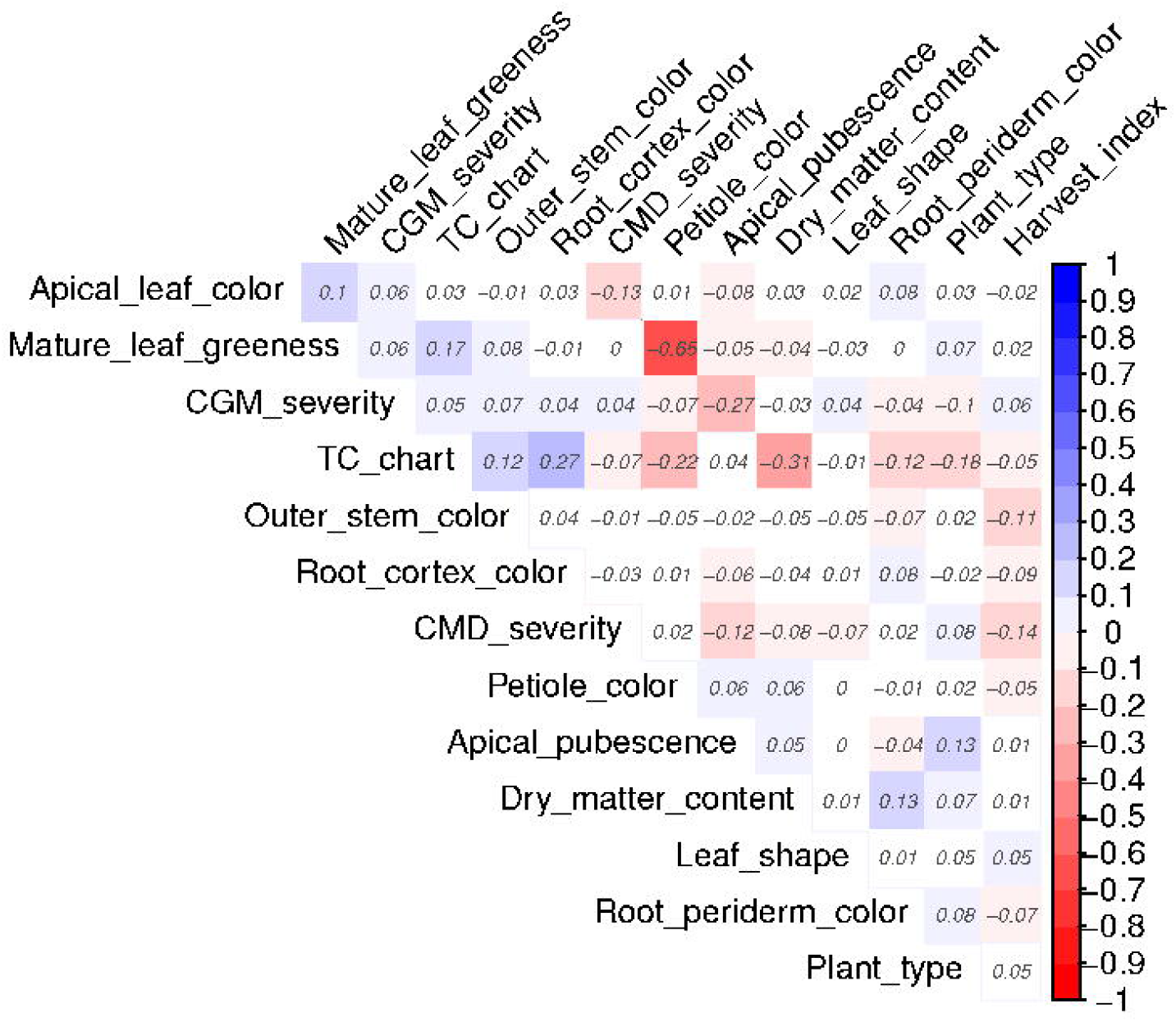
Heatmap of pairwise trait correlations using BLUPS from the 14 traits

### Distribution of SNP Markers and Population stratification

Genotyping of the panel enabled the identification of 202,789 imputed SNP markers. Upon filtering markers with minor allele frequency less than 1% we retained 101,521 SNPs. All the 101K SNPs were mapped onto the 18 chromosomes covering a total of 532.5 Mb of the cassava genome and a SNP density of 5.6 variants/Kbp (Figure 2). Individually, the SNP coverage per chromosome ranged from 3,821 on chromosome 7 to 11,189 on chromosome 1. The average minor allele frequency was 0.196 and 80% of the SNPs had frequency greater than 5% indicating enrichment of common SNP alleles in the population (Figure 2). To estimate the mapping resolution for our panel, we assessed pairwise linkage disequilibrium (LD) between 101,521 SNPs across the 5,130 cassava accessions. We used the mean r2 value as an estimate of LD decay using a window of 1 Mbp, followed by fitting a non-linear regression curve of LD versus distance. The whole-genome LD decay peaked at r2 of 0.349 and dropped to an r2 of 0.212 at a distance of 10 Kb (Figure 2).

**Figure 2.**
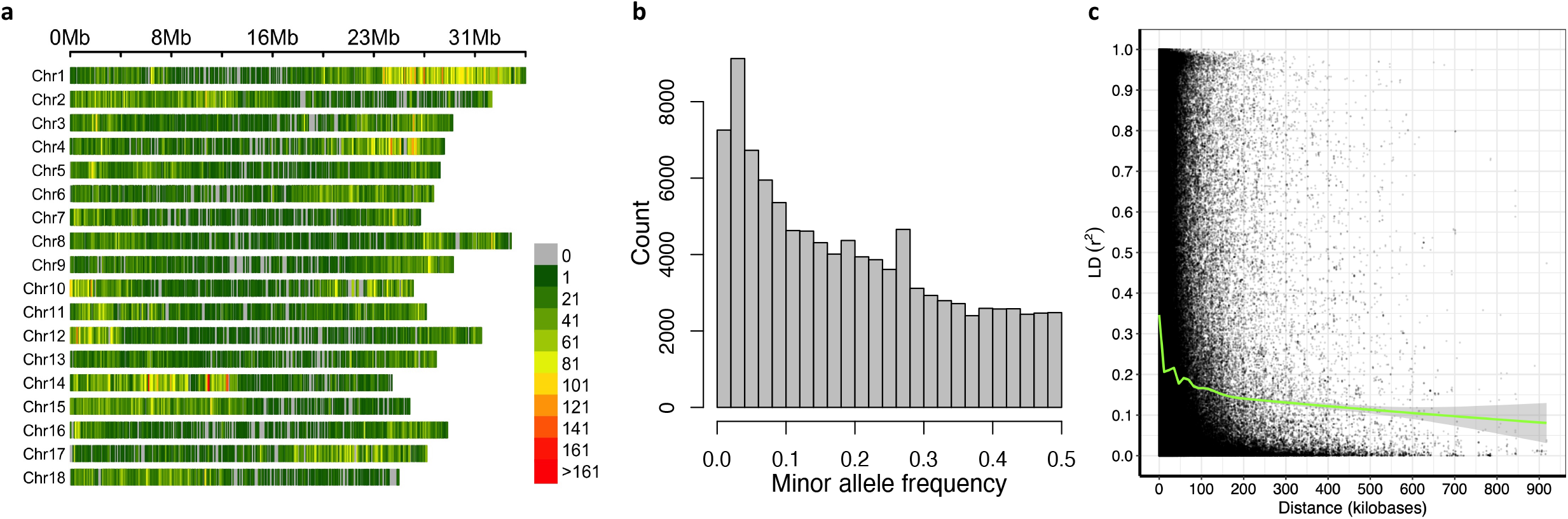
Overview of SNP genotyping data. (a) The density of SNPs on the 18 chromosomes of the cassava association mapping panel within 100 Kb window. (b) Histogram of minor allele frequency distribution. (c) Genome-wide Linkage disequilibrium (LD) decay for the cassava accessions in the panel showing the squared correlations (r2) between markers by marker physical distance (kb). The blue smoothening curve (LOESS) and the average LD were fitted to the LD decay.

Principal component analysis (PCA) was conducted to visualize the extent and degree of population structure present within the panel. While the first two principal components (PC) explained 7.4% of the total phenotypic variation, PC3 to PC10 together explained about 14% of the total variation (Figure 3). We observed a considerable overlap between cycles (C0, C1 and C2); hence, no distinct clusters were detected in the panel. The results of the identity-by-state distance showed a considerable range of relationship in the association panel with an average value of 0.23 and ranged from 0.02 to 0.32. We observed familial relationships along the diagonal with a few large blocks of closely related individuals. The off-diagonal indicated low kinship (Figure 3).

**Figure 3.**
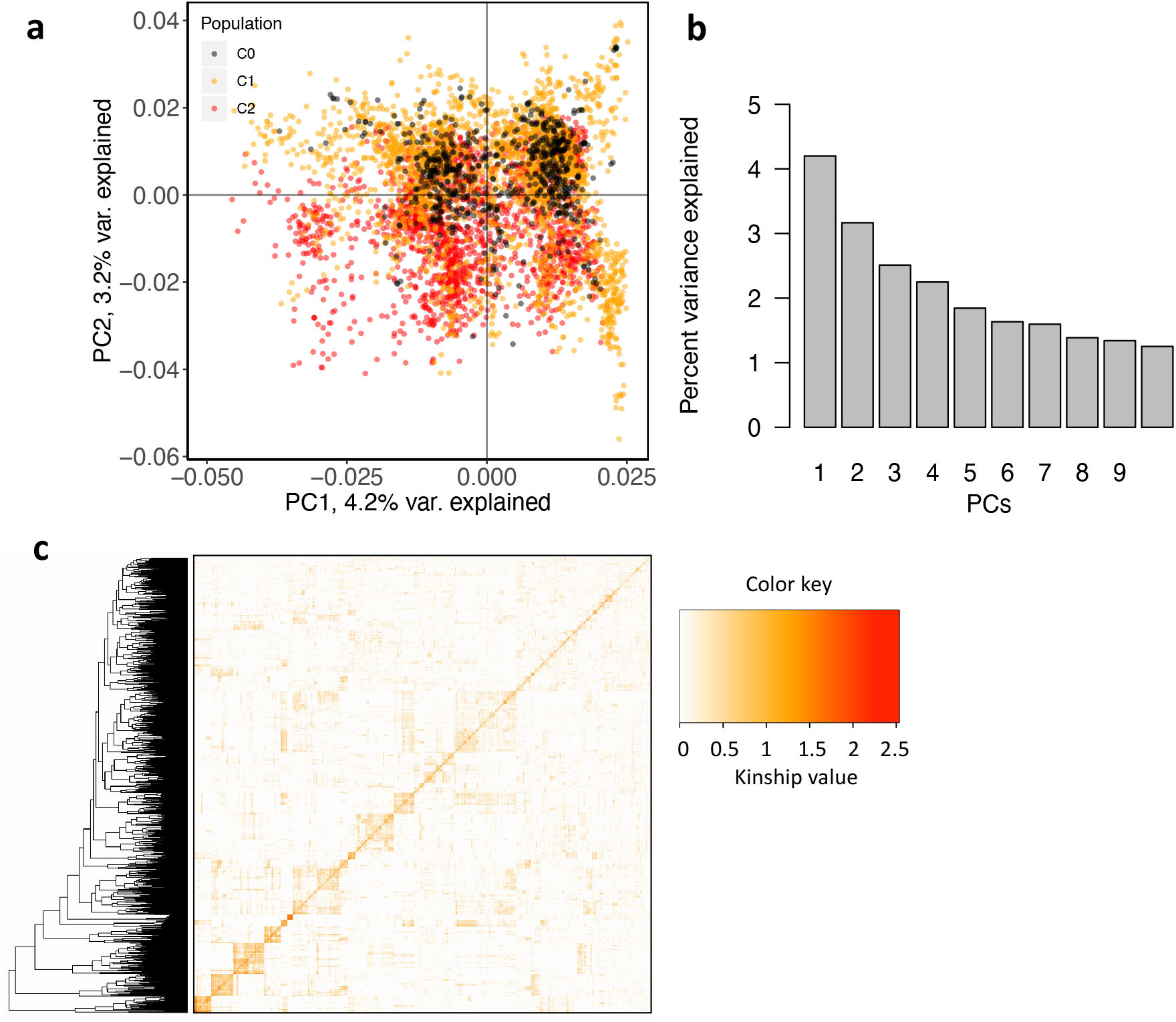
An assessment of population structure based on Principal Components Analysis of 101,521 SNP marker data (MAF>1%) for 5,130 individual cassava clones. (a) Scatter plot of the first two PCs; (b) the proportion of genetic variation explained by the first ten 10 PCs; and (c) Heatmap showing pairwise Kinship matrix

### Genome-wide association results

We identified a total of 27 unique genomic regions significantly associated with variation in the 14 studied traits following the MLMi analysis (Figure 4). Additional loci were uncovered for a majority of the traits when we considered the MLMe approach, bringing the total number of loci to 41 (Supplementary Figure S2). The most significant trait-marker associations from MLMi and MLMe for each trait and genomic region combinations are also provided in Table 3. In the following sections, we present the results and provide discussion for each trait starting with economically important traits to morphological traits.

**Figure 4.**
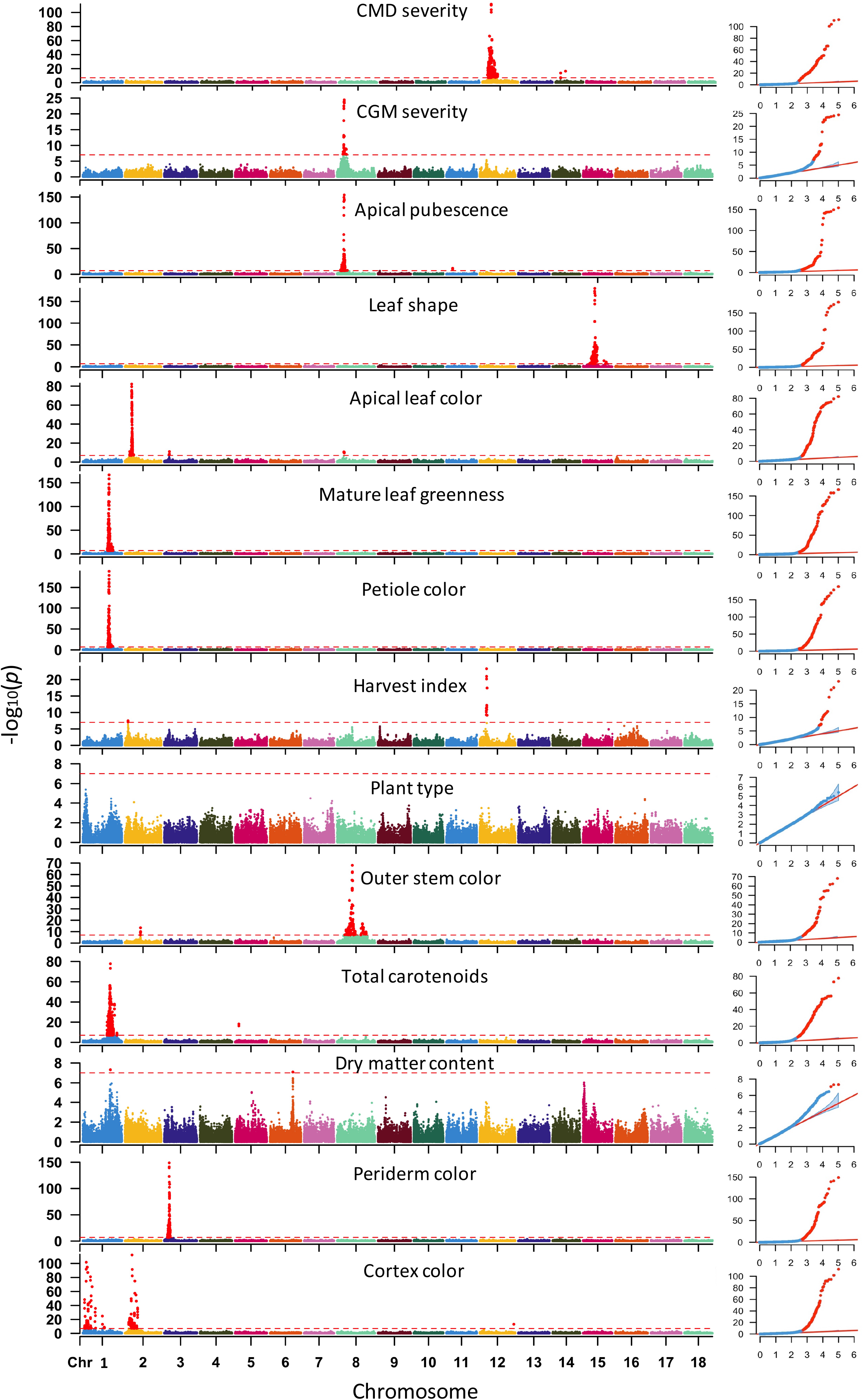
Manhattan plots for GWAS for 14 traits of 5,130 cassava accessions using MLM analysis approach. A total of 101,521 SNP markers were used for the GWAS analyses with the red horizontal line representing Bonferroni adjusted genome-wide significance threshold (α=0.05/101521=4.93 × 10^-7^). The QQ-plots inset - right with observed p-values on the y-axis and expected p-values on the x-axis

**Table 3.** Summary of most significant SNP markers at each major trait linked locus for the 14 studied traits

| Trait | SNP | Minor Allele | Major Allele | MAF | beta | se | P-value | beta* | se* | P-value* |
| --- | --- | --- | --- | --- | --- | --- | --- | --- | --- | --- |
| CMD severity | S12_7926132 | G | T | 0.44 | 0.82 | 0.04 | **1.7×10<sup>-112</sup>** | 0.89 | 0.02 | **P ≈ 0.0** |
| CMD severity | S14_4626854 | A | G | 0.40 | -0.23 | 0.03 | **1.7×10<sup>-14</sup>** | -0.23 | 0.03 | **1.0×10<sup>-14</sup>** |
| CMD severity | S14_9004550 | T | C | 0.17 | 0.28 | 0.03 | **4.2×10<sup>-17</sup>** | 0.27 | 0.03 | **1.7×10<sup>-16</sup>** |
| CGM severity | S1_28672656 | A | T | 0.28 | 0.09 | 0.04 | 1.4×10<sup>-2</sup> | 0.10 | 0.02 | **1.1×10<sup>-10</sup>** |
| CGM severity | S8_6409580 | C | G | 0.41 | -0.18 | 0.02 | **3.6×10<sup>-25</sup>** | -0.18 | 0.01 | **4.7×10<sup>-37</sup>** |
| CGM severity | S12_5524524 | C | T | 0.38 | -0.07 | 0.02 | 6.3×10<sup>-5</sup> | -0.07 | 0.01 | **1.3×10<sup>-7</sup>** |
| CGM severity | S17_23749968 | A | G | 0.31 | 0.09 | 0.02 | 1.5×10<sup>-5</sup> | 0.08 | 0.01 | **1.1×10<sup>-7</sup>** |
| Apical pubescence | S8_6409580 | C | G | 0.41 | 0.41 | 0.02 | **1.1×10<sup>-154</sup>** | 0.42 | 0.01 | **4.3×10<sup>-293</sup>** |
| Apical pubescence | S9_1588034 | A | G | 0.47 | 0.08 | 0.02 | **4.2×10<sup>-7</sup>** | 0.08 | 0.01 | **2.0×10<sup>-14</sup>** |
| Apical pubescence | S11_5727254 | A | G | 0.09 | -0.19 | 0.03 | **1.9×10<sup>-12</sup>** | -0.19 | 0.03 | **1.8×10<sup>-12</sup>** |
| Apical pubescence | S12_5524524 | C | T | 0.38 | 0.07 | 0.02 | 2.3×10<sup>-4</sup> | 0.09 | 0.01 | **6.8×10<sup>-18</sup>** |
| Apical pubescence | S16_1501762 | G | A | 0.30 | -0.06 | 0.02 | 2.8×10<sup>-3</sup> | -0.06 | 0.01 | **8.1×10<sup>-8</sup>** |
| Leaf shape | S15_10273255 | A | G | 0.03 | 2.70 | 0.10 | **3.7×10<sup>-174</sup>** | 2.90 | 0.08 | **P ≈ 0.0** |
| Leaf shape | S15_20573383 | G | C | 0.02 | 0.73 | 0.11 | **1.8×10<sup>-11</sup>** | 1.47 | 0.10 | **1.6×10<sup>-52</sup>** |
| Apical leaf colour | S2_6086714 | A | T | 0.43 | -1.22 | 0.06 | **6.1×10<sup>-83</sup>** | -1.26 | 0.04 | **5.7×10<sup>-267</sup>** |
| Apical leaf colour | S3_4745233 | G | A | 0.23 | -0.35 | 0.06 | **4.4×10<sup>-9</sup>** | -0.34 | 0.04 | **1.7×10<sup>-18</sup>** |
| Apical leaf colour | S8_6061421 | G | C | 0.41 | -0.32 | 0.05 | **1.9×10<sup>-11</sup>** | -0.27 | 0.03 | **2.4×10<sup>-16</sup>** |
| Mature leaf greenness | S1_23452638 | T | A | 0.16 | -1.11 | 0.04 | **2.7×10<sup>-167</sup>** | -1.24 | 0.03 | **P ≈ 0.0** |
| Petiole colour | S1_23452638 | T | A | 0.16 | 2.73 | 0.10 | **9.8×10<sup>-180</sup>** | 2.73 | 0.10 | **9.8×10<sup>-180</sup>** |
| Harvest index | S2_2809137 | G | T | 0.09 | -0.04 | 0.01 | **3.0×10<sup>-8</sup>** | -0.04 | 0.01 | **5.6×10<sup>-15</sup>** |
| Harvest index | S12_6055806 | A | G | 0.24 | -0.03 | 0.00 | **5.4×10<sup>-24</sup>** | -0.02 | 0.00 | **2.0×10<sup>-17</sup>** |
| Plant type | S1_3192405 | T | C | 0.28 | 0.20 | 0.05 | 1.7×10<sup>-5</sup> | 0.22 | 0.04 | **3.8×10<sup>-8</sup>** |
| Plant type | S1_25303195 | C | A | 0.48 | 0.23 | 0.05 | 3.4×10<sup>-5</sup> | 0.27 | 0.04 | **5.3×10<sup>-14</sup>** |
| Outer stem colour | S2_13928566 | C | G | 0.39 | -0.44 | 0.10 | **3.6×10<sup>-14</sup>** | -0.45 | 0.06 | **6.5×10<sup>-15</sup>** |
| Outer stem colour | S8_13604799 | G | A | 0.34 | -1.06 | 0.06 | **8.3×10<sup>-69</sup>** | -1.06 | 0.05 | **8.7×10<sup>-89</sup>** |
| Outer stem colour | S8_22630799 | G | A | 0.25 | -0.58 | 0.07 | 3.3×10<sup>-17</sup> | -0.76 | 0.06 | **1.3×10<sup>-37</sup>** |
| Total carotenoids (color chart) | S1_24159583 | T | C | 0.30 | 0.27 | 0.02 | **5.3×10<sup>-57</sup>** | 0.37 | 0.01 | **1.3×10<sup>-219</sup>** |
| Total carotenoids (color chart) | S1_24636113 | G | A | 0.23 | 0.49 | 0.03 | **1.8×10<sup>-78</sup>** | 0.57 | 0.02 | **1.3×10<sup>-270</sup>** |
| Total carotenoids (color chart) | S1_30543962 | G | A | 0.10 | 0.18 | 0.03 | **6.3×10<sup>-8</sup>** | 0.40 | 0.02 | **2.4×10<sup>-76</sup>** |
| Total carotenoids (color chart) | S5_3387558 | T | C | 0.10 | 0.21 | 0.02 | **5.5×10<sup>-17</sup>** | 0.20 | 0.02 | **2.0×10<sup>-16</sup>** |
| Total carotenoids (color chart) | S8_4319215 | A | C | 0.01 | 0.22 | 0.05 | 5.2×10<sup>-5</sup> | 0.23 | 0.04 | **3.4×10<sup>-8</sup>** |
| Total carotenoids (color chart) | S8_25598183 | T | G | 0.03 | 0.18 | 0.04 | 6.3×10<sup>-6</sup> | 0.18 | 0.03 | **1.1×10<sup>-8</sup>** |
| Total carotenoids (color chart) | S15_7659426 | G | T | 0.26 | -0.07 | 0.02 | 2.0×10<sup>-3</sup> | -0.06 | 0.01 | **3.2×10<sup>-7</sup>** |
| Total carotenoids (color chart) | S16_484011 | G | T | 0.35 | 0.09 | 0.03 | 7.6×10<sup>-4</sup> | 0.05 | 0.01 | **8.9×10<sup>-8</sup>** |
| Dry matter content | S1_24636113 | G | A | 0.23 | -1.32 | 0.24 | **5.0×10<sup>-8</sup>** | -1.68 | 0.14 | **1.7×10<sup>-33</sup>** |
| Dry matter content | S6_20589894 | G | A | 0.48 | 0.85 | 0.17 | **3.4×10<sup>-7</sup>** | 0.78 | 0.09 | **1.7×10<sup>-16</sup>** |
| Dry matter content | S12_5524524 | C | T | 0.37 | 0.68 | 0.17 | 9.8×10<sup>-5</sup> | 0.69 | 0.10 | **8.0×10<sup>-12</sup>** |
| Dry matter content | S15_1012346 | C | T | 0.44 | -0.95 | 0.20 | 2.2×10<sup>-6</sup> | -0.84 | 0.10 | **4.0×10<sup>-17</sup>** |
| Dry matter content | S16_25663808 | T | C | 0.35 | -0.48 | 0.18 | 6.8×10<sup>-3</sup> | -0.69 | 0.10 | **4.2×10<sup>-12</sup>** |
| Periderm colour | S3_4545411 | G | C | 0.43 | -0.38 | 0.02 | **1.4×10<sup>-123</sup>** | -0.42 | 0.01 | **7.3×10<sup>-288</sup>** |
| Cortex colour | S1_3047840 | T | G | 0.01 | 1.08 | 0.05 | **1.6×10<sup>-92</sup>** | 1.18 | 0.05 | **1.8×10<sup>-135</sup>** |
| Cortex colour | S2_6566608 | C | T | 0.01 | 0.81 | 0.04 | **5.0×10<sup>-83</sup>** | 0.89 | 0.04 | **2.0×10<sup>-113</sup>** |

A1 = Reference allele (the default effect allele), A2 = Alternate allele, MAF = Frequency of the reference allele, B = SNP effect, SE = standard error of SNP effect, P = marker-trait association p-value. Bold fonts represent markers that are significant at the Bonferroni threshold of $0.05/101,521 = 4.93 \times 10^{-7}$
\* SNP effect, standard error and p-values obtained from GCTA MLMe model.

### Cassava mosaic disease severity

The genetic basis of CMD resistance has been studied extensively using bi-parental linkage mapping (Rabbi et al. 2014) and GWAS (Wolfe et al. 2016). These studies repeatedly showed that the main resistance to the disease is conferred by a single major gene on chromosome 12 which is widely known as CMD2 locus (Akano et al. 2002). In the present study, we uncovered the same locus and two other loci on chromosome 14. The CMD2 region on chromosome 12 is tagged by marker S12_7926132 (p-value = 1.7 ×10^-112^). The favourable allele at this marker (T) was found to occur at high frequency in the population (Freq > 0.66). The two additional loci on chromosome 14 are tagged by markers S14_4626854 (p-value = 1.7 × 10^-14^) and S14_9004550 (p-value = 4.2 × 10^-17^). The SNP at the CMD2 locus had a larger effect (β value 0.82) compared to the two new loci on chromosome 14 (β value of 0.23 and 0.28).

### Cassava green mite severity

We found four genomic regions associated with CGM resistance in our panel (Figure 4) of which only marker S8_6409580, occurring around 6.41 Mb region of chromosome 8, was previously reported (Ezenwaka et al. 2018). The remaining loci were found on chromosomes 1, 12 and 17 (Table 3). We note that except for the markers on chromosome 8, which were significant in both MLMi and MLMe, the remaining loci were only significant in the MLMe analysis, though their SNP effects (β statistic) were very similar in the two analyses. Association analysis of the related trait, apical pubescence identified significant loci on five chromosomes, two of which co-locate with same regions underlying resistance to CGM on chromosomes 8 (S8_6409580) and 12 (S12_5524524). The genetic correlation between the two traits is −0.51 in the population (Supplementary Table S2). The other loci were on chromosomes 9 (S9_1588034), 11 (S11_5727254), and 16 (S16_1501762).

### Carotenoid content

GWAS analysis of colour-chart based variation in root carotenoid content revealed a major locus on chromosome 1 tagged by three markers around 24.1, 24.6 and 30.5 Mb regions with the top markers being S1_24159583, S1_24636113 and S1_30543962, respectively. Previous GWAS analyses reported significant associations in this region (Esuma et al. 2016; Rabbi et al. 2017; Ikeogu et al. 2019). In addition, we uncovered five new genomic regions associated with this trait on chromosome 5 (around 3.38 Mb), 8 (two peaks at 4.31 and 25.59 Mb regions), and 15 (7.65 Mb), and 16 (0.48 Mb). We note that the regions in the last three chromosomes, including two on chromosome 8, were only detected via the MLMe analysis.

### Dry matter content

GWAS for variation in dry matter content following the MLMi analysis revealed two major loci for DM of which one was previously reported (Rabbi et al. 2017). The most significant locus occurs on chromosome 1 around 24.64 Mb region and is tagged by marker S1_24636113 (p-value = 5.0 × 10^-8^). The second locus was tagged by marker S6_20589894 (p-value = 3.4 × 10^-7^). Additional loci on chromosomes 12 (S12_5524524, p-value = 8.0 × 10^-12^), 15 (S15_1012346, p-value = 4.0 × 10^-17^), and 16 (S16_25663808, p-value = 4.2 × 10^-12^) were detected from the MLMe analysis. Along with the new locus on chromosome 6, these regions have not been previously reported to be associated with dry matter content variation and are suitable for further genetic studies including identification of underlying candidate genes.

### Harvest index

The MLMi-based GWAS for harvest index, the ratio of fresh root weight to total plant weight, uncovered two genomic regions that are significantly associated with the trait. The first peak is in chromosome 2, tagged by SNP S2_2809137 (p-value of p-value = 3 × 10^-8^). The second locus occurred on chromosome 12 with SNP S12_6055806 showing the strongest association with the trait (p-value = 5.4 × 10^-24^). Analysis of the same trait using the MLMe approach uncovered several other regions scattered across chromosomes 3, 4, 6, 8, 9, 14, 15, 16, and 18.

### Morphological traits

GWAS for outer cortex colour uncovered two association signals located on chromosomes 1 (3.05 Mbp region) and 2 (6.56 Mbp region) and tagged by SNPs S1_3047840 (p-value 1.6 × 10^-92^) and S2_6566608 (p-value 5.0 × 10^-83^), respectively. A single genomic region on chromosome 3 (4.54 Mbp) was found to be linked to periderm colour. The most significant SNP at this locus (S3_4545411) had a p-value of 1.4 × 10^-123^.

Two association signals were detected for plant type from MLMe while no marker passed the Bonferroni threshold in the MLMi analysis. The detected loci jointly occurred on chromosome 1 at 2.19 Mbp region (tagged by SNP S1_3192405, p-value 3.82 × 10^-8^) and 25.30 Mbp region (S1_25303195, 5.25 × 10^-14^). Three loci were found to be associated with stem colour variation, one in chromosome 2 and two on chromosome 8. The most significant locus was around 13.6 Mbp region of chromosome 8 and is tagged by marker S8_13604799 (p-value 8.3 × 10^-69^). The other two loci were tagged by SNPs S2_13928566 (p-value 3.6 ×10^-14^) and S8_22630799 (p-value 3.3 × 10^-17^).

Genetic architecture of leaf morphology traits showed that they are controlled by one to three major loci, indicating simple genetic architecture. We found a single genomic region around 23.45 Mbp of chromosome 1 to be associated with this trait and is tagged by SNP S1_23452638 (p-value 9.8 × 10^-180^). Notably, the same exact SNP was found to underlie the variation in mature leaf greenness. It is therefore not surprising that these traits are negatively correlated in our population. SNP effect analysis showed that while allele “T” at S1_23452638 had a positive effect on petiole colour, the same allele showed a negative effect on leaf greenness. Regression of the marker on the traits for leaf colour and petiole colour returned an R2 0.57 and 0.62, respectively.

Variation in the colour apical leaves was found to be associated with 3 loci occurring on chromosomes 2, 3, and 8. The most significant marker was S2_6086714 (p-value 6.1 × 10^-83^) followed by the markers on chromosome 8 (S8_6061421, p-value 1.9 × 10^-11^) and 3 (S3_4745233, p-value 4.4 × 10^-9^). Multiple regression returned an R^2^ of 0.31 for this trait suggesting either a more complex architecture or imprecise scoring of the variation in the trait. The GWAS result for leaf shape uncovered two major loci on chromosome 15 occurring around 10.27 Mbp and 20.57 Mbp regions. The first peak tagged by SNP S15_10273255 was highly significant (3.7 × 10^-174^) while the second peak was tagged by S15_20573383 (p-value 1.8 × 10^-11^). Fitting linear model with the two top SNPs for this trait returned R^2^ of 0.40.

### Genetic architecture of studied traits

To assess the genetic architecture of the studied traits, we fit a linear model using peak SNPs at the identified loci (Table 3) as independent variables against the traits BLUPS as the response variables. Peak SNPs at the identified loci explained approximately 34% of the trait variation on average and R2 ranged from 5% to 62% (Figure 5). Markers for cassava green mite severity, harvest index, plant type and dry matter content had the lowest predictive ability (R^2^ < 0.1). Most of the morphology and colour related traits for leaves and roots had between 1 and 3 peaks of association except apical leaf pubescence with 6 loci. Peaks associated with variation in total carotenoid content and resistance to CMD had a large effect (R^2^ = 0.60 and 0.45, respectively). The major loci controlling these traits had known candidate genes reported previously. Still, the new loci identified in these as well as the other traits are attractive candidates for follow-up studies.

**Figure 5.**
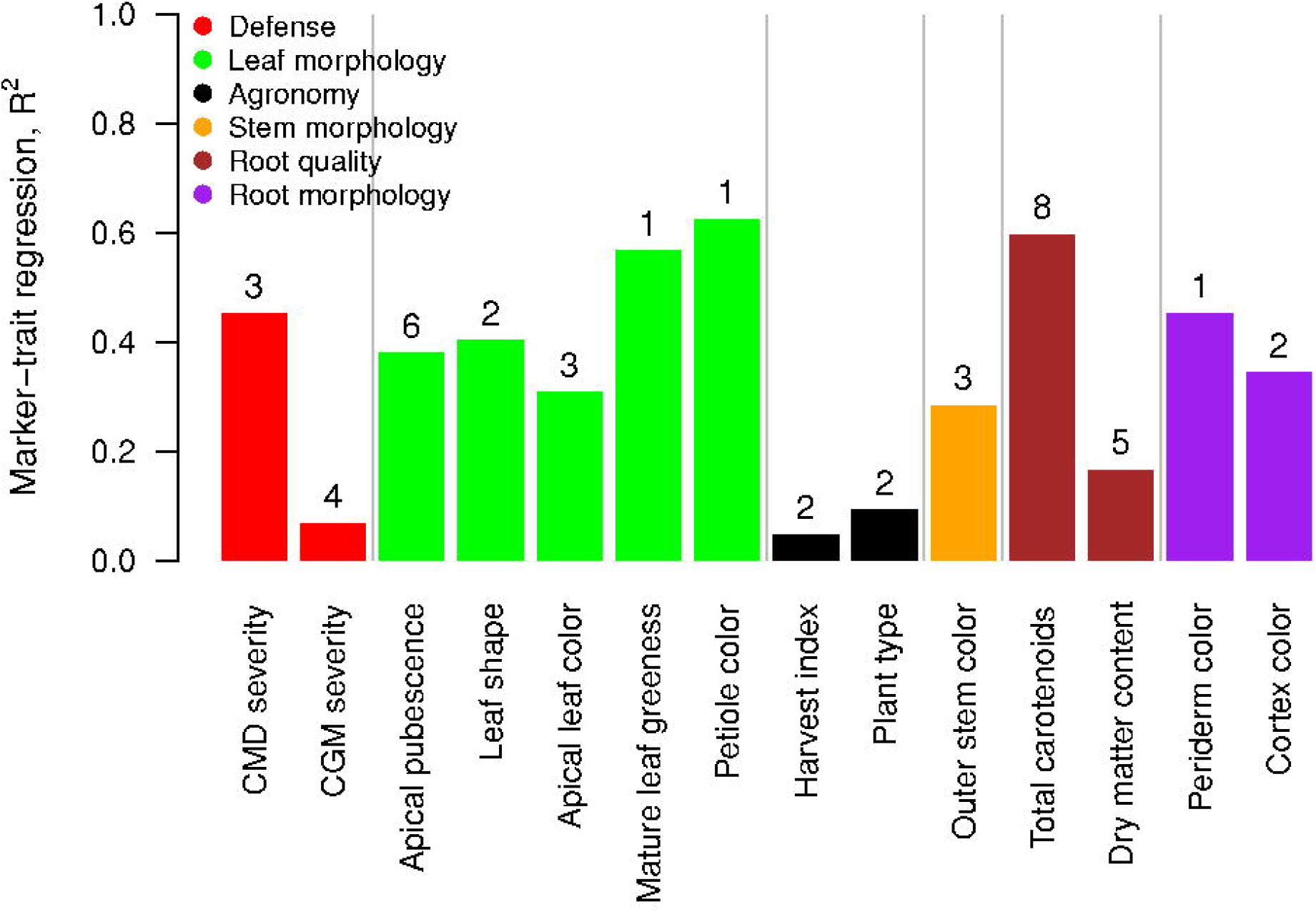
Multiple Marker trait regression barplot across traits and the proportion of phenotypic variance explained. Numbers above bar plot denote the number of loci in the regression model

### Identification of favorable alleles

To identify favourable alleles for traits under selection in the breeding pipeline, the most significantly associated SNP (lowest p-value) at each major-effect locus were chosen. Allelic substitution effects for these markers are shown in Supplementary Figure S3. Selected traits include resistance to CMD and CGM, increased dry matter and carotenoid content. For CMD resistance, the haplotypes at the top SNPs S12_7926132 (allele G/T), S14_4626854 (A/G) and S14_9004550 (T/C) are *T-A-C*, respectively. There were 251 genotypes that were fixed for the favourable alleles in the population. Their average CMD severity BLUP was the lowest among all haplotype combinations (mean = −0.56, SD = 0.29). We also note dominance allele effect at the first SNP which is linked to the CMD2 locus, which agrees with previous results from biparental QTL studies (Akano et al. 2002; Rabbi et al. 2014).

The CGM resistance-linked haplotype for SNPs S1_28672656 (A/T), S8_6409580 (C/G), S12_5524524 (C/T) and S17_23749968 (A/G) are *T-C-C-G*, respectively. Alleles associated with increased pubescence particularly for loci that co-locates with CGM resistance of chromosomes 8 (S8_6409580 (C/G)) and 12 (S12_5524524 (C/T)) are *C-T*.

Several SNP loci from chromosome 1 (S1_24159583, S1_24636113, S1_30543962), 5 (S5_3387558), 8 (S8_4319215, S8_25598183), 15 (S15_7659426), and 16 (S16_484011) were associated with increased carotenoid content. The favourable haplotype for chromosome 1 SNPs are *T-G-G*, respectively while that for chromosome 5 is *T*. The favourable haplotype for the two loci on chromosome 8 is *A-T* while for chromosomes 15 and 16 SNPs it is *T-T*, respectively. No individuals in the population were fixed for the favourable alleles across all the SNPs.

The favourable allele at the most significant dry matter locus on chromosome 1 (S1_24636113 allele G/A) is *A*. We note that this marker also co-located with the major locus for carotenoid content and allele *A* has a non-favourable effect in carotenoid content. The favourable haplotypes at the other loci on chromosomes 6 (S6_20589894 G/A), 12 (S12_5524524, C/T), 15 (S15_1012346 C/T) and 16 (S16_25663808 T/C) are *G-C-T-C*. A total of 61 genotypes fixed for the favourable alleles at the SNPs on chromosomes 1, 6 and 12. Their average dry matter BLUP was the highest among all haplotype combinations (mean = 2.62, SD = 3.66).

### Candidate gene identification

The most significant GWAS peaks were further investigated for the presence of potential candidate genes using local linkage disequilibrium, coupled to gene annotation. We matched the highly significant SNP markers identified in the MLMe analysis with gene annotation in the regions up and downstream derived from phytozome online database. Seventeen candidate genes that colocalized with the identified putative genomic loci on height chromosomes were retrieved from the cassava genome available on phytozome v.12.1 and are presented in Supplementary Table S3. Several of these identified genes were further selected and highlighted based on their biological significance within a given biological pathway. Additionally, we provide regional Manhattan plots for each locus-trait combination in Supplementary Figure S4. These plots include candidate genes within 100 Kb of the top SNP marker (50 kb upstream, 50 kb downstream) with some adjustments based on the extent of local linkage disequilibrium with the candidate SNP.

Three genes associated with total carotenoid content (TC-chart) were identified on chromosomal regions 1, 5 and 15. Of the three associated genes, Manes.01G124200 (Phytoene synthase) occurred within the previously reported genomic region (Esuma et al. 2016; Rabbi et al. 2017). Manes.01G124200 gene has a transferase enzymatic activity critical in the carotenoid biosynthesis pathway. The other two genes are novel: Manes.05G051700 and Manes.15G102000 (both of which are Beta-carotene dioxygenases) located at 3.87 Mb on chromosome 5 and 7.58 Mb on chromosome 15, respectively, and are also known to play major roles in carotenoid biosynthesis.

For dry matter content, we identified two genes involved in starch and sucrose metabolism occurring 600 Kbp away from top SNP S1_24636113 in chromosome 1: Manes.01G123000 (UDP-Glucose pyrophosphorylase) and Manes.01G123800 (Sucrose synthase) previously reported (Rabbi et al. 2017). Although these genes are a few hundred Kb away from the top SNP, this particular genomic region of chromosome 1 is known to harbour extensive LD in Africa cassava germplasm (Rabbi et al. 2017). Other candidate genes found close to the top SNPs on chromosomes 6, 15 and 16 are Manes.06G103600 (Bidirectional sugar transporter Sweet4-Related); Manes.15G011300 (Sweet17 homologue, which mediates fructose transport across the tonoplast of roots) and Manes.16G109200 (Hexokinase), respectively.

Our search for candidate genes related to CMD severity uncovered two peroxidase genes: Manes.12G076200 and Manes.12G076300 occurring less than 45 Kbp away from marker S12_7926132 in chromosome 12. These two candidate genes were previously reported by Wolfe et. al. (2016) and Rabbi et al. (2014). Peroxidases have been reported to play a role in activating plant defence systems upon pathogen infections (Ye et al. 1990; Wu et al. 1997; Gonçalves et al. 2013).

Our analyses further detected three loci that colocalized with three candidate genes associated with CGM severity. Specifically, SNP marker S8_6409580 fell in the coding region of Manes.08G058000 gene, a homolog of AtMYB16, encoding a MIXTA-like MYB gene which regulates cuticle development and trichome branching (Oshima et al. 2013). Structural traits including trichomes and waxy cuticles are known to act as a physical barrier to arthropod pest attachment, feeding and oviposition (Mitchell et al. 2016)

Harvest index GWAS analysis highlighted two regions on chromosome 2 and 12, respectively. On chromosome 2, the candidate SNP (S2_2809137) is located 24 Kbp away Manes.02G035900 a homologue of BFRUCT4 (vacuolar invertase) a key enzyme in sucrose hydrolysis and involved in the export of reduced carbon sink organs such as roots (Haouazine-Takvorian et al. 1997; Nägele et al. 2010). For leaf shape, a candidate gene Manes.15G136200, homologous to KNOX1 that is implicated in the expression of diverse leaf shapes in plants was found around 186 Kb away from the major locus on chromosome 15 tagged by SNP S15_10273255 (Furumizu et al. 2015). Candidate gene search around the major locus for petiole colour and leaf greenness tagged by SNP S1_23452638 revealed the presence of a Myb transcription factor homologue Manes.01G115400 which occurred 30.7 Kb away from the top marker. Myb genes are known to play a key role in regulating pigment biosynthesis pathway in plants (Nesi et al. 2001; Kobayashi et al. 2002; Himi and Noda 2005; Allan et al. 2008; Furumizu et al. 2015)

## Discussion

Understanding the genetic architecture of key breeding-goal traits is a critical step towards more efficient and accelerated genetic improvement. This study builds on and extends previous cassava GWAS efforts by analysing a large breeding population phenotyped extensively over successive years and stages of selection in multi-environment field trials. The population showed large phenotypic variation among clones with respect to all traits (Supplementary Figure S1). Furthermore, the population was derived from two successive cycles of recurrent selection using elite parents with good breeding values for yield, dry matter content and resistance against CMD (Wolfe et al. 2016). The collection is therefore expected to be enriched for favourable alleles from major- and minor-effect loci underlying these traits and is therefore well suited to efficiently conduct a marker-trait association study.

The observed and consistent trend in the magnitude of both broad-sense and SNP-based heritability estimates further emphasizes the significant contribution of additive genetic factors in the expression of some of these traits. The heritability estimates recorded in our study also give an indication of good repeatability and reproducibility of the experimental procedures. The heritability estimates we found are comparable to those previously reported in other studies for these traits (Oliveira et al. 2014; Oliveira et al. 2015; Njoku et al. 2015; Silva et al. 2016; Favour et al. 2017; Rao et al. 2018).

Accounting for population structure and genetic relatedness in GWAS is necessary to reduce false positives. PCA did not reveal the presence of substantial population stratification in our GWAS panel. This is not surprising since extensive inter-crosses are routinely carried out as part of the generation of new genetic variation in the IITA Cassava Breeding program. Moreover, individuals from C1 and C2 largely overlapped with each other and also with the founder population (GG). GWAS analysis using PCA and the kinship matrix gave very similar results to the analysis that considered kinship alone in controlling for spurious associations. For this reason, as well as for computational efficiency, we used the MLM model for the full analysis.

Our data replicated the previously identified associations for CMD and CGM resistance traits, dry matter and carotenoid content, and also showed stronger evidence of major gene effect in this larger sample. In addition to the confirmed loci, we uncovered additional genomic regions with significant associations that were previously not reported. For example, two additional regions on chromosome 14 were found to contribute to increased resistance to CMD virus and present useful targets for further genetic analysis. Genotypes carrying favourable resistance alleles at these as well as the CMD2 major effect locus on chromosome 12 exhibit high levels of resistance against CMD and rarely show symptoms. However, despite the concerted global efforts to identify the causal gene underlying the CMD2 locus in cassava, there has not been any breakthrough in cloning and functionally identifying the actual gene. We hope that these resources will guide the cassava community in narrowing down on the candidate genes to carry out the functional analysis.

Although our main interest was in traits that are under active breeding selection, we also measured a number of morphological traits that we knew to be heritable. Among the morphological traits presented here, to our best knowledge, only leaf shape and apical leaf pubescence were previously reported (Ezenwaka et al. 2018; Zhang et al. 2018). While our study confirmed major locus for apical pubescence locus on chromosome 8 we did not replicate the results of Zhang et al. (2018) for leaf aspect ratio suggesting a possibility of different genetic factors.

Many traits of interest to crop improvement are positively or negatively correlated. Such correlations can cause unfavourable changes in traits that are important but that are not under direct selection. Alternatively, one trait can be used to indirectly select for another positively correlated trait which is more difficult to phenotype. Genetic correlations among traits can arise due to linkage disequilibrium or pleiotropy (a single gene having multiple otherwise unrelated biological effects, or shared regulation of multiple genes) (Chen and Lübberstedt 2010). Correlations due to linkage disequilibrium tend to be temporary and are generally considered to be less important than pleiotropy. Of particular importance is the negative correlation observed between total carotenoid content variation and dry matter content which confirms previous findings (Ceballos et al. 2013; Njoku et al. 2015; Esuma et al. 2016; Ceballos et al. 2017; Rabbi et al. 2017; Okeke et al. 2018). Both genetic linkage and pleiotropy are plausible reasons for the inverse relationship. The linkage basis is supported by the co-location of major QTLs for these traits on chromosome 1 and the presence of major genes in the biochemical pathways in close proximity. On the other hand, genetically engineered cassava to produce and accumulate carotenoids in storage roots was found in one study to have reduced dry matter content which indicates a pleiotropic effect (Beyene et al. 2017). Besides the region on chromosome 1, we identified additional loci on chromosomes 5, 8, 15, and 16 that additively explain more than 60% of the phenotypic variation for total carotenoids. These and other minor effect loci could explain the lack of inverse relationship between carotenoids and dry matter content in other populations (Ceballos et al. 2013). Likewise, we also found several significant regions associated with dry matter content although they only explain 16% of the trait variation. Other notable negative but the favourable correlation in the study population was between apical pubescence and CGM severity. Generally, genotypes with glabrous apical leaves are more affected by the pest than pubescent ones (Raji et al. 2008; Chalwe et al. 2015; Ezenwaka et al. 2018). For both traits, we identified a common major locus on chromosome 8 that is associated with variation in the degree of pubescence as well as CGM damage severity. We also found other loci that were unique to either trait suggesting additional factors may be contributing to the resistance against CGM besides the presence of trichomes. Significant genetic and phenotypic correlations were not observed between harvest index, dry matter content and plant type, implying that they are amenable to concurrent improvement.

### Opportunities and implications

The present study is based predominantly on germplasm developed by the IITA breeding program and therefore represents a subset of the available diversity worldwide. While some traits such as yield and yield components are universally considered, other traits, especially biotic and abiotic stresses, are region-specific. Moreover, restricted germplasm movement due to quarantine regulations makes it nearly impossible to evaluate the same population in multiple regions simultaneously. Further studies using germplasm from other regions including east Africa and Latin America – the centre of origin and diversity of cassava – is expected to reveal region and/or population-specific large effect alleles. Such efforts are expected to enrich the catalogue of major effect loci available for molecular breeding.

A major objective of this study was to provide breeders with a catalogue of major loci for marker-assisted selection. Many of the previous QTL studies using bi-parental mapping populations in cassava have had limited value due to low marker densities and poor genetic resolution (Ferguson et al. 2012; Ceballos et al. 2015; Hershey 2017). Access to high-density genome-wide SNP markers through GBS coupled with GWAS mapping approach has resulted in higher mapping resolution, to within a few Kb known candidate genes for several traits in the present study. If converted to allele-specific high-throughput SNP assays, SNPs tagging major loci from the present study can be used to screen and identify individuals carrying favourable alleles during early stages of selection. However, further validation of these loci is required to ensure they are effective across environments (genotype-by-environment interaction) and populations (different genetic background) before large-scale deployment. For other traits such as harvest index, the identified loci only contributed to a small proportion of trait variation suggesting additional genes with small effects and thus are more likely to benefit from genomic selection (GS) to estimate breeding values from genome-wide marker data. GS relies on high-density marker coverage such that each QTL is likely to be in linkage disequilibrium (LD) with at least one marker (Goddard and Hayes 2007). A tandem approach incorporating MAS and GS is likely to increase breeding efficiency, reduce breeding cycle and cost (Zhao et al. 2014). MAS can be used to screen large number of individuals at seedling nursery stages to cull accessions that do not carry favourable alleles. Reduced number of lines can then be genotyped at higher density for GS and allocated to field testing plots for evaluation and selection for complex traits.

## Conclusions

In this study, we demonstrate that the use of a diverse association mapping panel consisting of landraces and improved cultivars from multiple breeding initiatives can identify SNP variants associated with agronomically essential traits in cassava. The power of this study to discover additional markers associated with measured traits was derived from extensive multi-locational testing over four years at four IITA testing locations in Nigeria. The SNP markers we identified provide a useful reference catalogue not only for cassava breeding programs but also studies aimed at uncovering unique biological pathways necessary for advancing genetic transformation studies. To realize the full potential of these marker-trait associations in population improvement, a critical next step is to validate these loci in independent populations before their deployment for routine use in marker-assisted selection.

## Acknowledgements

This work was funded by The Bill & Melinda Gates Foundation and the Department for International Development of the United Kingdom (award number OPP1048542) through the “Next Generation Cassava Breeding Project”; and the CGIAR-Research Program on Roots, Tubers and Bananas. Thanks to the technical staff at IITA Cassava Breeding Program for implementing the field experiments.

## Authors contributions

IYR, CE, JLJ, and PK conceived and designed the study; IYR, SIK, GB, AA, and MY performed analyses and wrote the manuscript; CE, EL, EP, MW, JLJ, and PK edited the manuscript; CA, KO, RU, ASI, and PP Implemented field trials, generated and curated data; and PK Provided overall coordination and leadership

## Competing interests

The authors declare no competing interests.

## Data availability

The data that support the findings of this study are openly available in CassavaBase at ftp://ftp.cassavabase.org/manuscripts/PlantMolBiol_Rabbi_et_al_2020/

**Supplementary Tables and Figures**

**Supplementary Table S1** Clone names, GBS ID, pedigree, clonal generation and BLUPS for the 14 traits.

**Supplementary Table S2** The pairwise estimates of genetic (lower diagonal) and phenotypic correlations (upper diagonal) using GBLUPS and BLUPS respectively among the 14 traits.

**Supplementary Table S3** Summary information of the potential candidate genes identified in the vicinity of the GWAS hits for analyzed traits.

**Supplementary Figure S1** Variation and trends in phenotypic data for 14 morphological, agronomic, quality and defensive traits in a diverse cassava association mapping panel evaluated between 2013 to 2016 across four locations. Histograms of 14 traits were based on de-regressed BLUPs distributions measured on 5,130 cassava accessions.

**Supplementary Figure S2** Manhattan plots for GWAS for 14 traits of 5,130 cassava accessions using the leave-one-chromosome-out “MLMe” analysis approach. A total of 101,521 SNP markers were used for that GWAS analyses with the red horizontal line representing genome-wide significance threshold (5%). The QQ-plots inset - right with observed p-values on the y-axis and expected p-values on the x-axis.

**Supplementary Figure S3** Allelic substitution effects in the most significant SNP at each locus identified for each of the 14 studied traits. Trait BLUPs distribution on y-axis and SNP genotype status of the marker on x-axis. SNPs were converted to dosage format, where 0, 1, and 2 indicates the copies of the minor alleles. The first allele in the suffix of a SNP name denotes the allele being counted in the dosage coding. For example, dosage score of 2 in SNP S12_7926132_G(T) means homozygous for “G”, a score of 1 means heterozygote “GT”, and a score of 0 means homozygote alternate allele “T”. (A) CMD severity; (B) CGM severity; (C) Apical leaf pubescence; (D) Leaf shape(E); Apical leaf colour; (F) Mature leaf greenness; (G) Petiole colour; (H) Harvest index; (I) Plant type; (J) Outer stem colour; (K) Total carotenoids content; (L) Dry matter content; (M) Periderm colour; (N) Root cortex colour.

**Supplementary Figure S4** Regional Manhattan plots for each locus-trait combination. Plots include candidate genes within 100 Kb of the top SNP marker (50 kb upstream, 50 kb downstream) with some adjustments based on the extent of local linkage disequilibrium with the candidate marker. SNPs are colour coded based on linkage disequilibrium with the top marker. (A) CMD severity; (B) CGM severity; (C) Apical leaf pubescence; (D) Leaf shape; (E) Apical leaf colour; (F) Mature leaf greenness; (G) Petiole colour; (H) Harvest index; (I) Plant type; (J) Outer stem colour; (K) Total carotenoids content; (L) Dry matter content; (M) Periderm colour; (N) Root cortex colour.

## Notes

### Competing Interest Statement

The authors have declared no competing interest.

