## Supplementary_tables_and _figs for "Genome-wide association analysis reveals new insights into the genetic architecture of defensive, agro-morphological and quality-related traits in cassava": ESM_SupplementaryFigures_news.pdf

**Electronic supplementary materials**

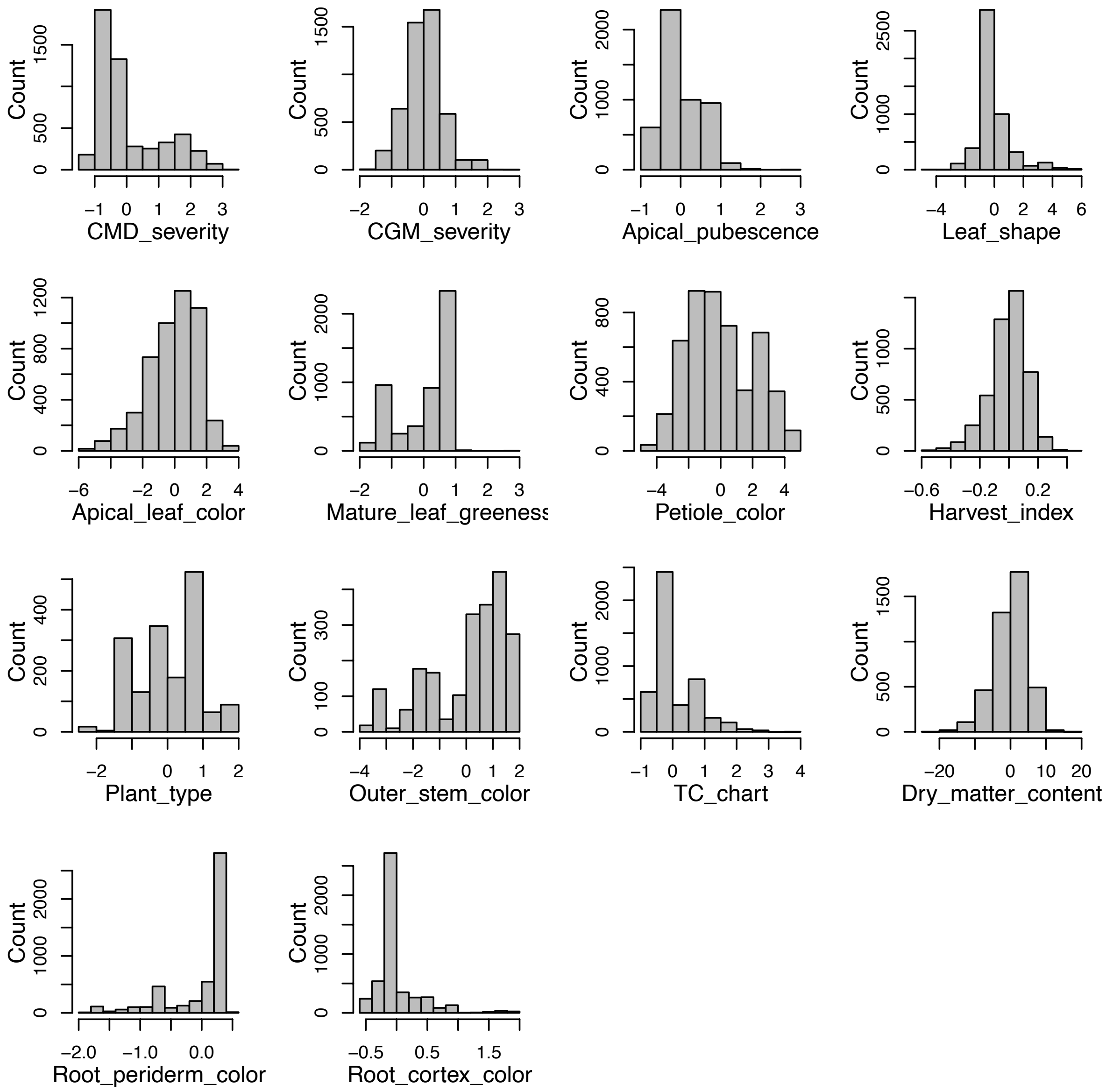


**Figure S1** Histogram showing the distribution of BLUPs for the 14 traits used in the GWAS analysis.

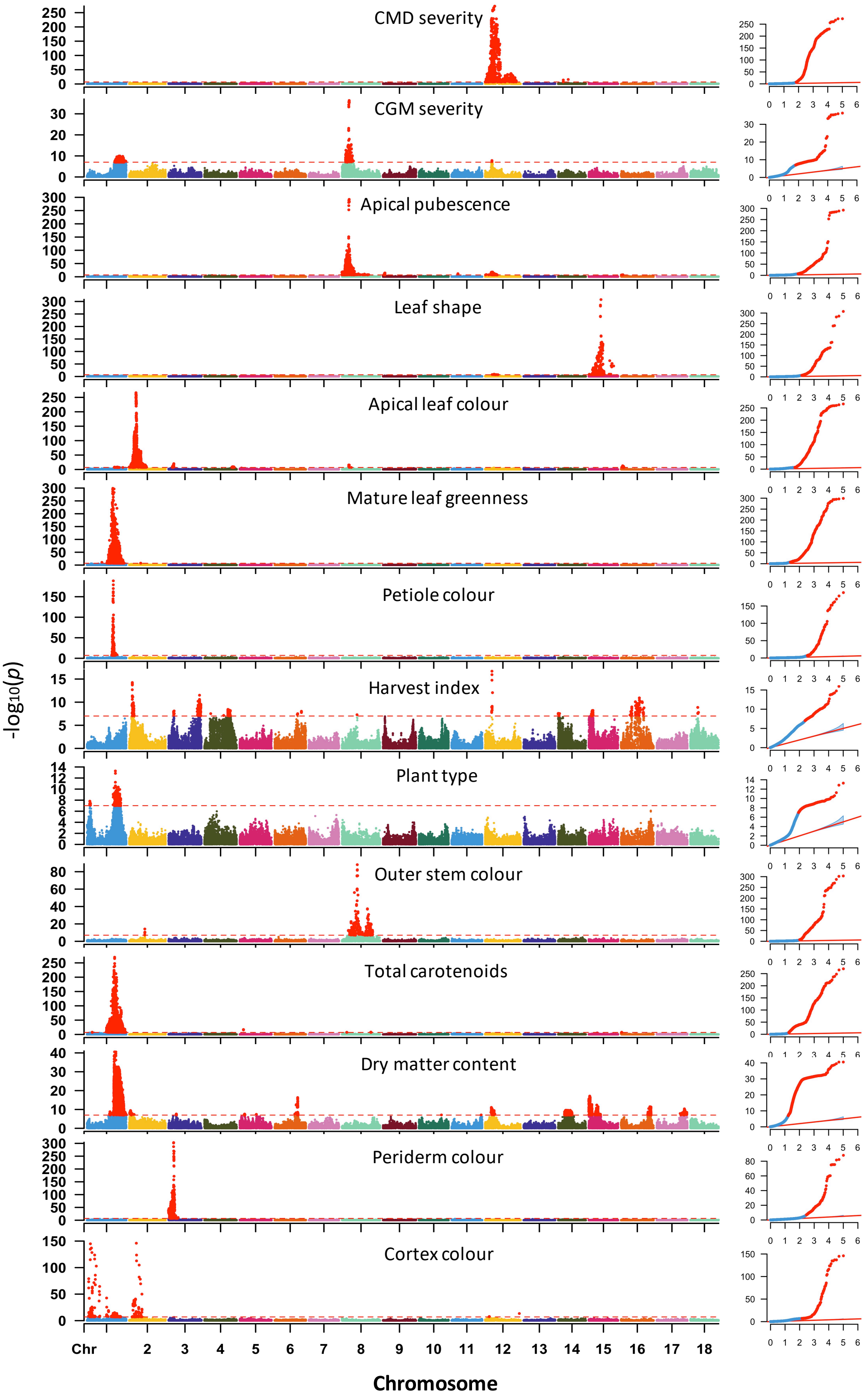


**Figure S2** Manhattan plots for GWAS for 14 traits of 5,130 cassava accessions using "MLMe" analysis approach. A total of 101,521 SNP markers were used for that GWAS analyses with the red horizontal line representing genome-wide significance threshold ($\alpha$=0.05/101521 = 4.93 × $10^{-7}$). The QQ-plots inset - right with observed p-values on the y-axis and expected p-values on the x-axis.

(A) CMD severity

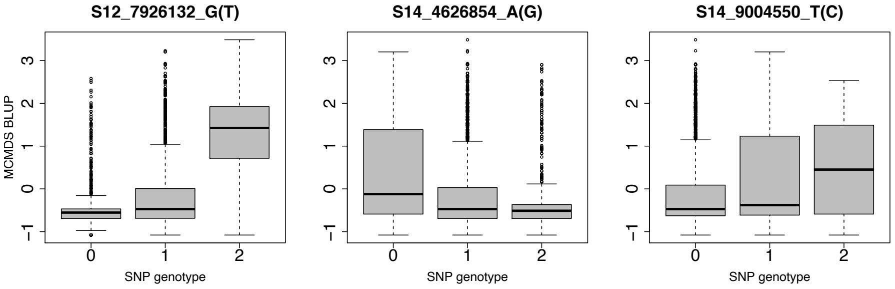


(B) CGM severity

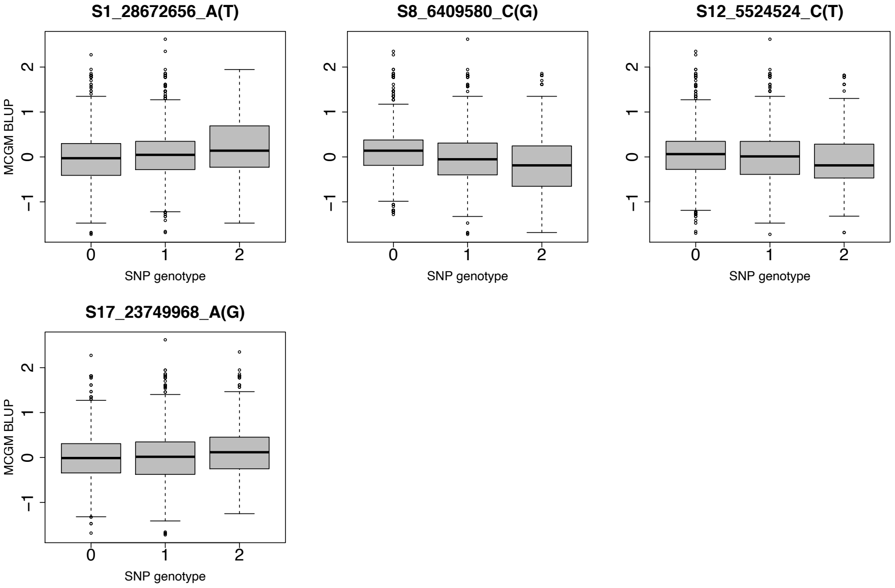


(C) Apical leaf pubescence

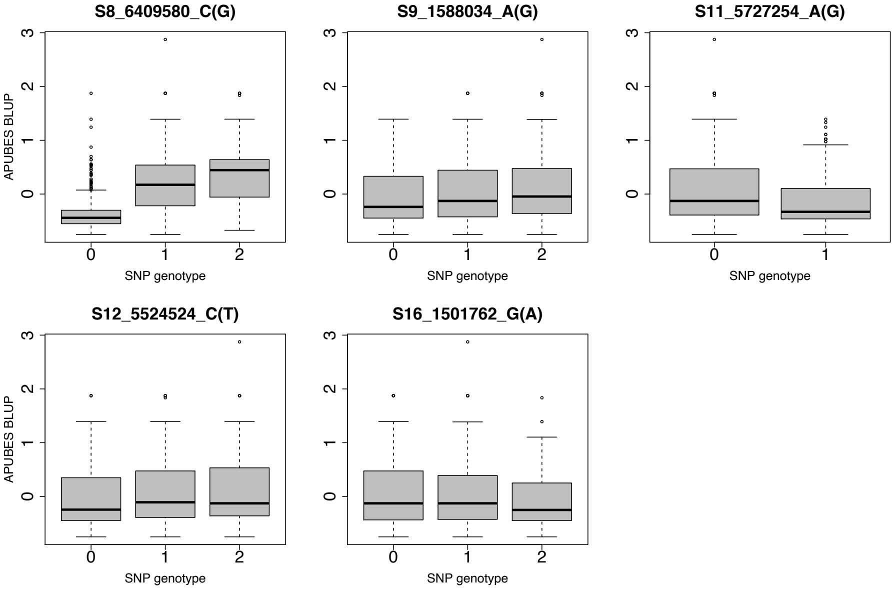

### (D) Leaf shape

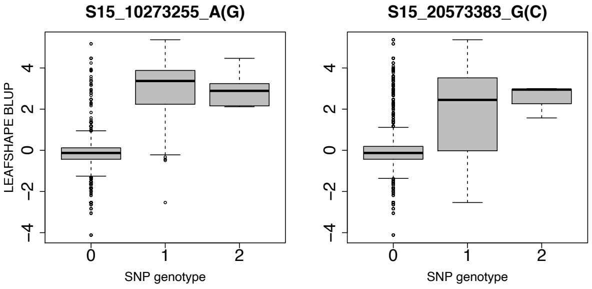


### (E) Apical leaf colour

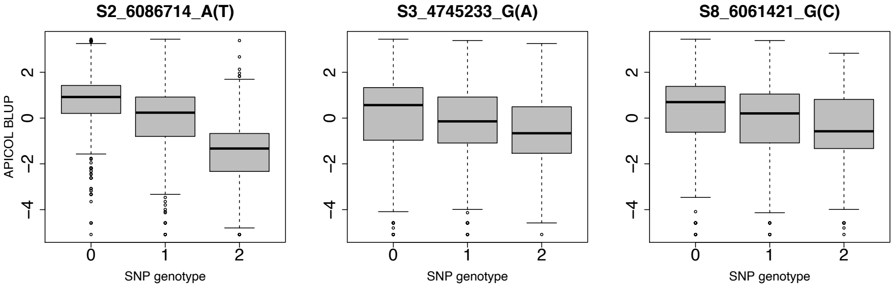


### (F) Mature leaf greenness

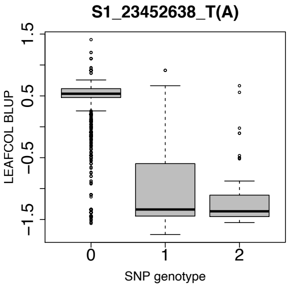


### (G) Petiole colour

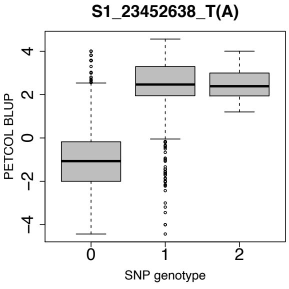


### (H) Harvest index

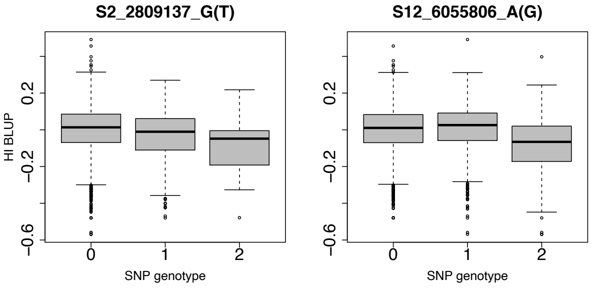

(I) Plant type

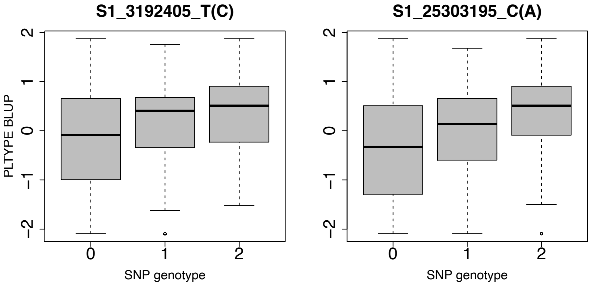


(J) Outer stem colour

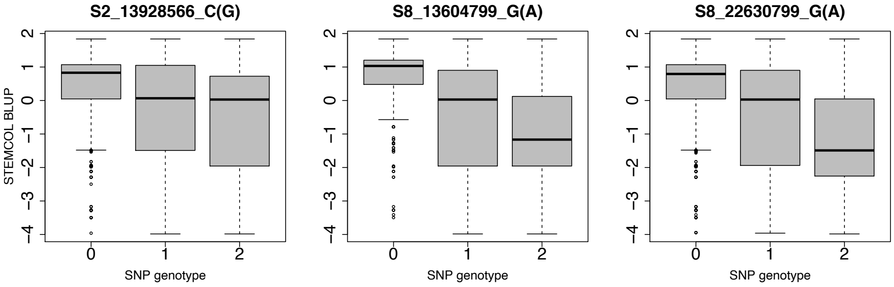


(K) Total carotenoids content

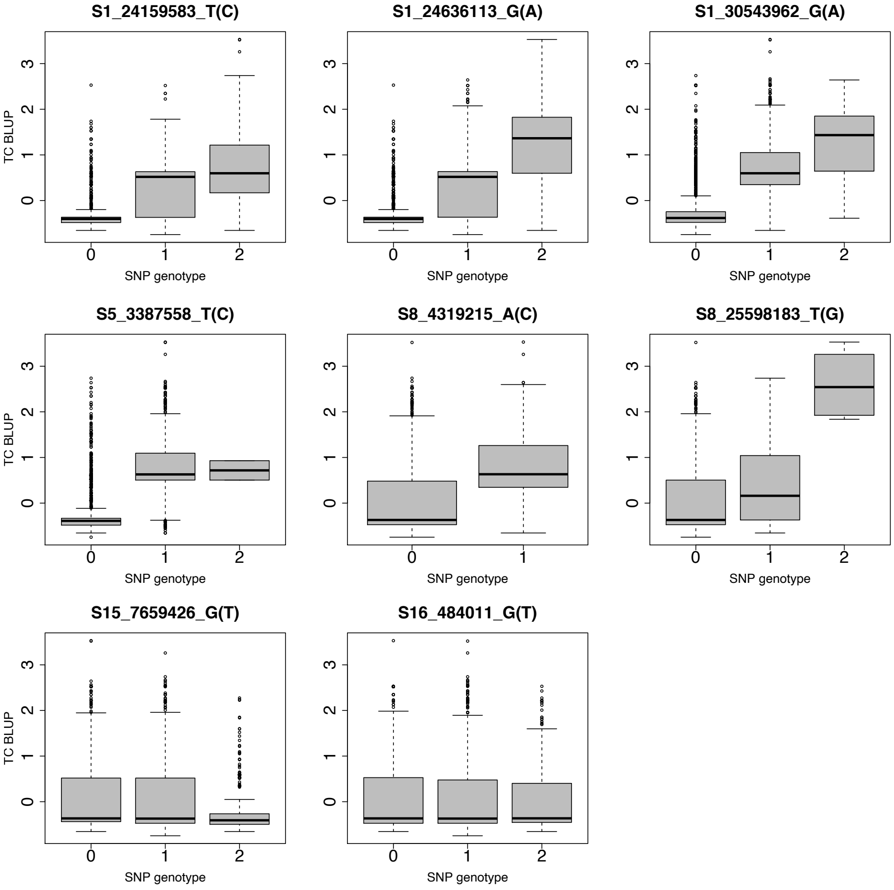

(L) Dry matter content

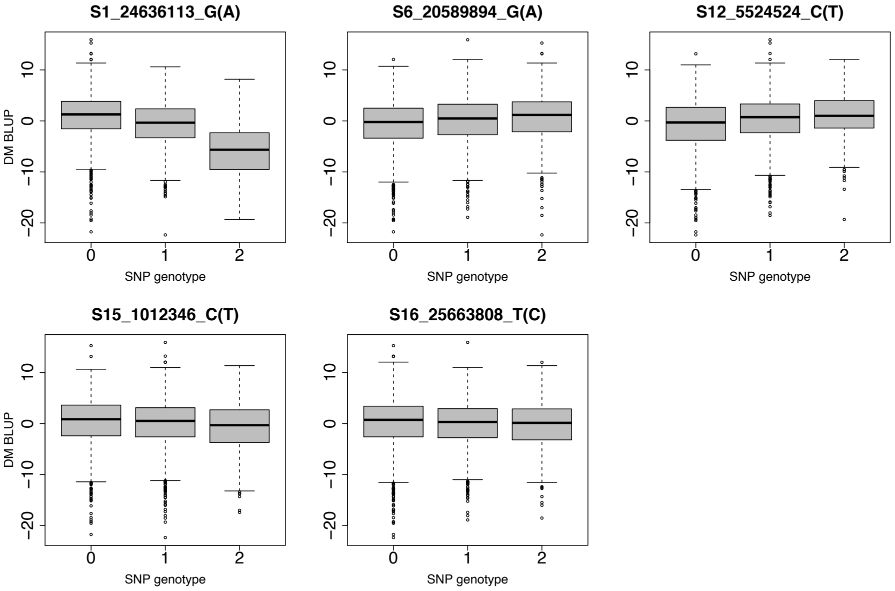


(M) Periderm colour

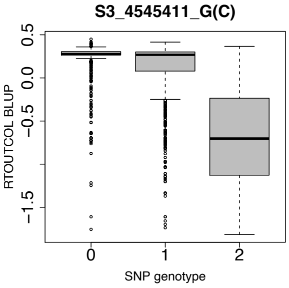


(N) Root cortex colour

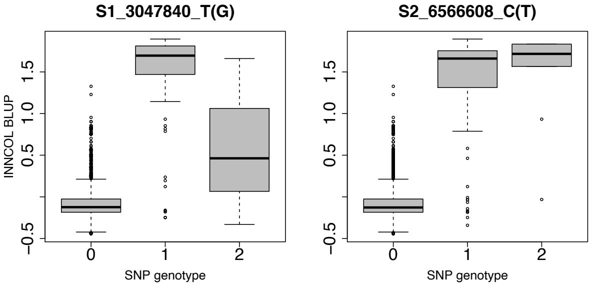


**Figure S3** Allelic substitution effects in the most significant SNP at each locus identified for each of the 14 studied traits. Trait BLUPs distribution on y-axis and SNP genotype status of the marker on x-axis. SNPs were converted to dosage format, where 0, 1, and 2 indicates the copies of the minor alleles. The first allele in the suffix of a SNP name denotes the allele being counted in the dosage coding. For example, dosage score of 2 in SNP S12_7926132_G(T) means homozygous for "G", a score of 1 means heterozygote "GT", and a score of 0 means homozygote alternate allele "T". (A) CMD severity; (B) CGM severity; (C) Apical leaf pubescence; (D) Leaf shape(E); Apical leaf colour; (F) Mature leaf greenness; (G) Petiole colour; (H) Harvest index; (I) Plant type; (J) Outer stem colour; (K) Total carotenoids content; (L) Dry matter content; (M) Periderm colour; (N) Root cortex colour.

**A**

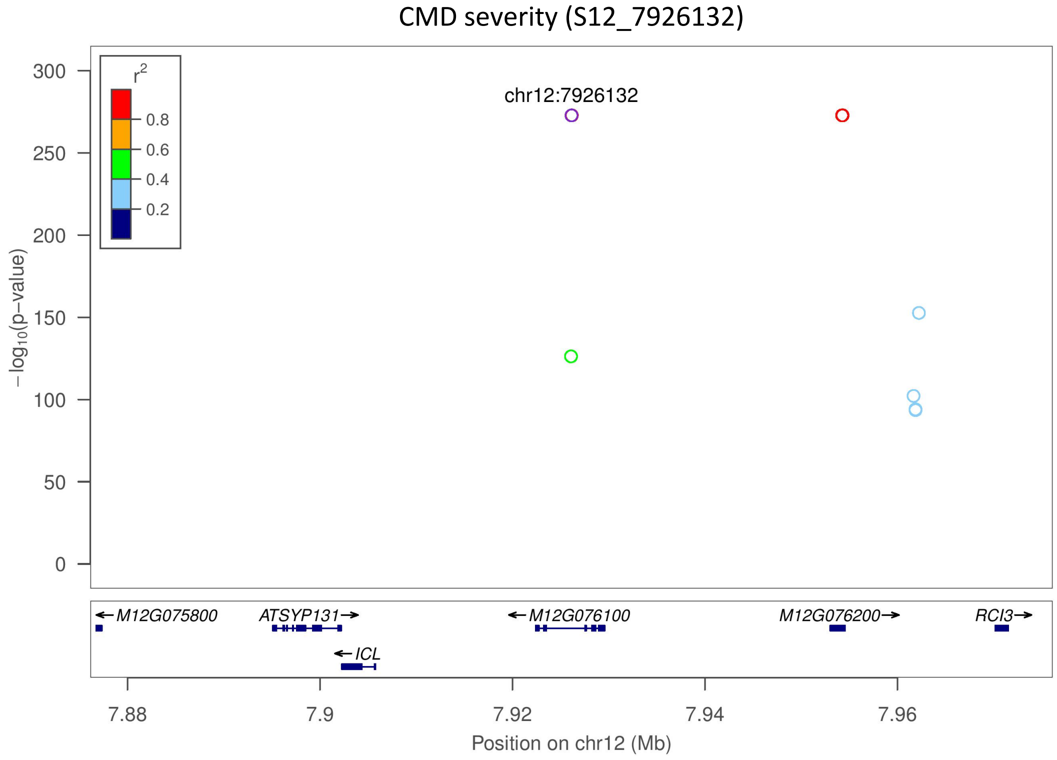
CMD severity (S12_7926132)
r2
0.8
0.6
0.4
0.2
chr12:7926132
−log10(p−value)
300
250
200
150
100
50
0
M12G075800
ATSYP131
ICL
M12G076100
M12G076200
RCI3
7.88
7.9
7.92
7.94
7.96
Position on chr12 (Mb)

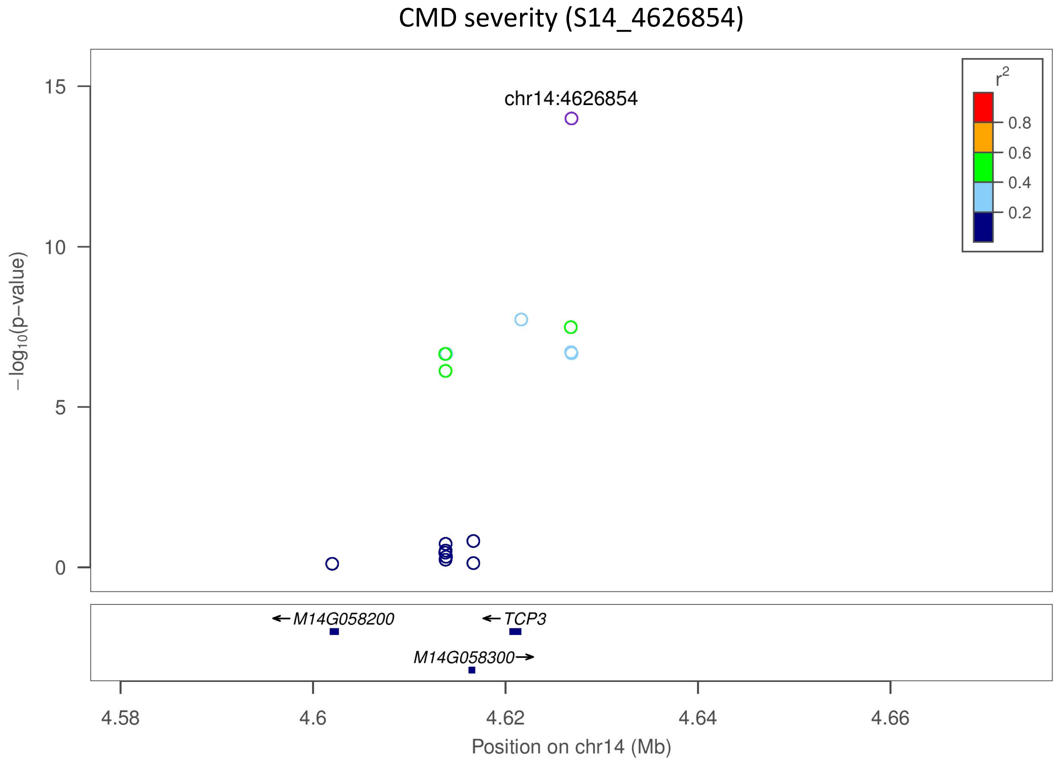
CMD severity (S14_4626854)
r2
0.8
0.6
0.4
0.2
chr14:4626854
−log10(p−value)
15
10
5
0
M14G058200
TCP3
M14G058300
4.58
4.6
4.62
4.64
4.66
Position on chr14 (Mb)

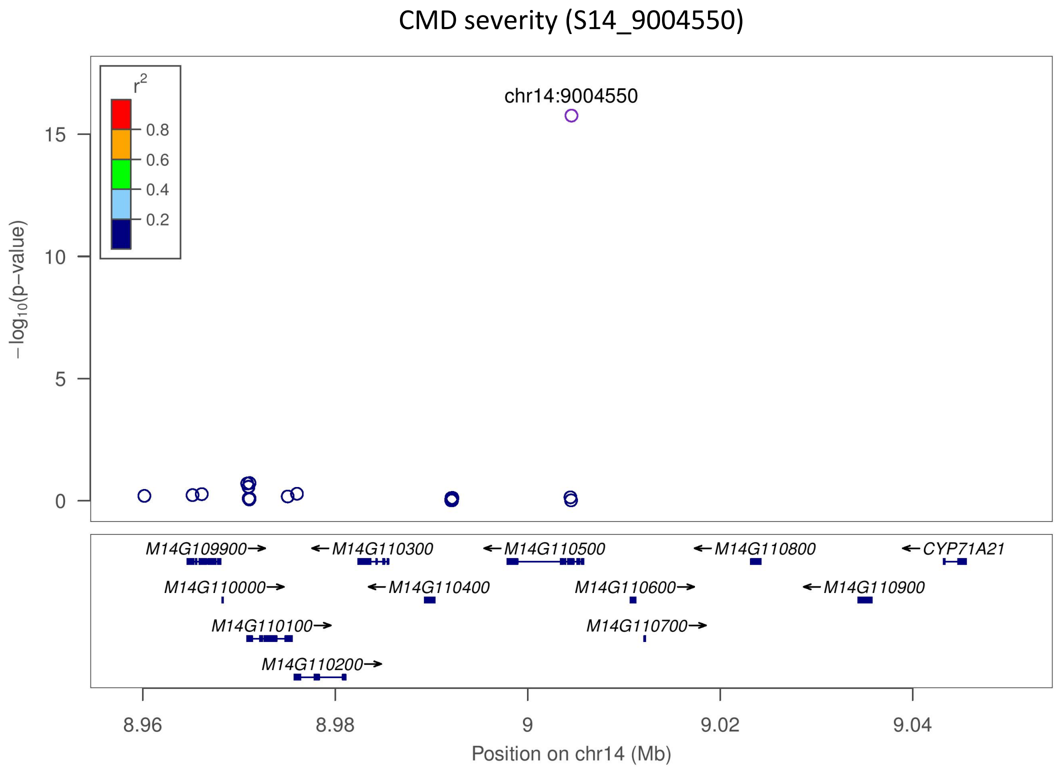
CMD severity (S14_9004550)
chr14:9004550
r2
0.8
0.6
0.4
0.2
−log10(p−value)
15
10
5
0
M14G109900→
←M14G110300
←M14G110500
←M14G110800
←CYP71A21
M14G110000→
←M14G110400
M14G110600→
←M14G110900
M14G110100→
M14G110700→
M14G110200→
8.96
8.98
9
9.02
9.04
Position on chr14 (Mb)

**B**

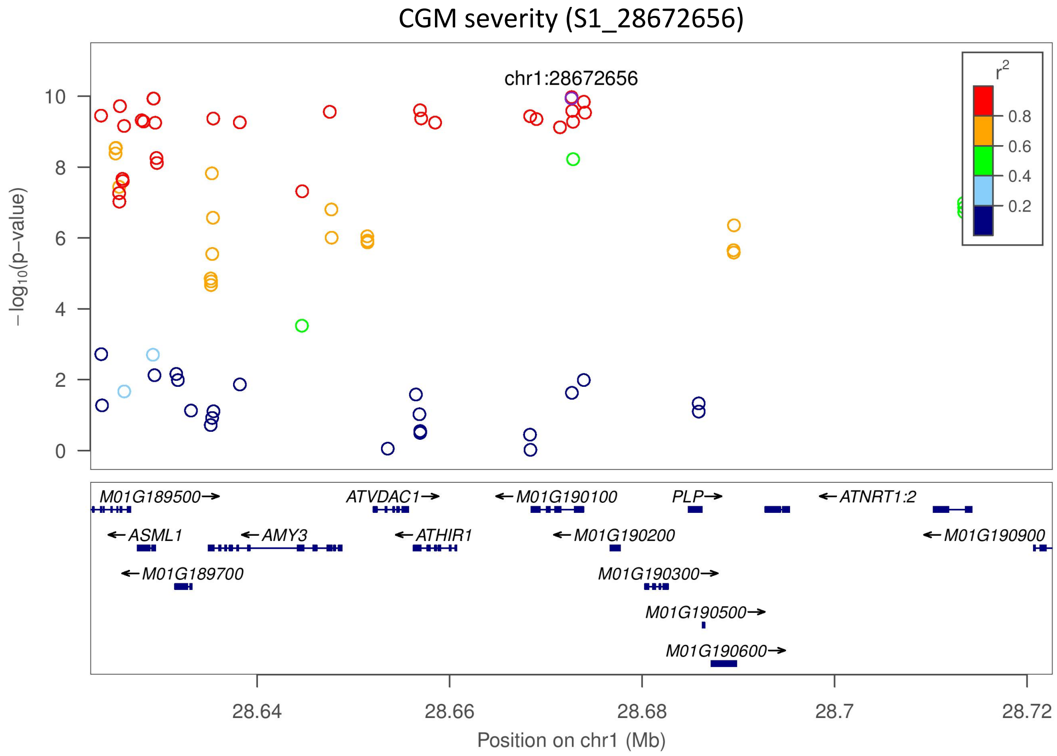
CGM severity (S1_28672656)
chr1:28672656
r2
0.8
0.6
0.4
0.2
−log10(p−value)
10
8
6
4
2
0
M01G189500→
ATVDAC1→
←M01G190100
PLP→
←ATNRT1:2
←ASML1
←AMY3
←ATHIR1
←M01G190200
←M01G190900
←M01G189700
M01G190300→
M01G190500→
M01G190600→
28.64
28.66
28.68
28.7
28.72
Position on chr1 (Mb)

CGM severity (S8_6409580)

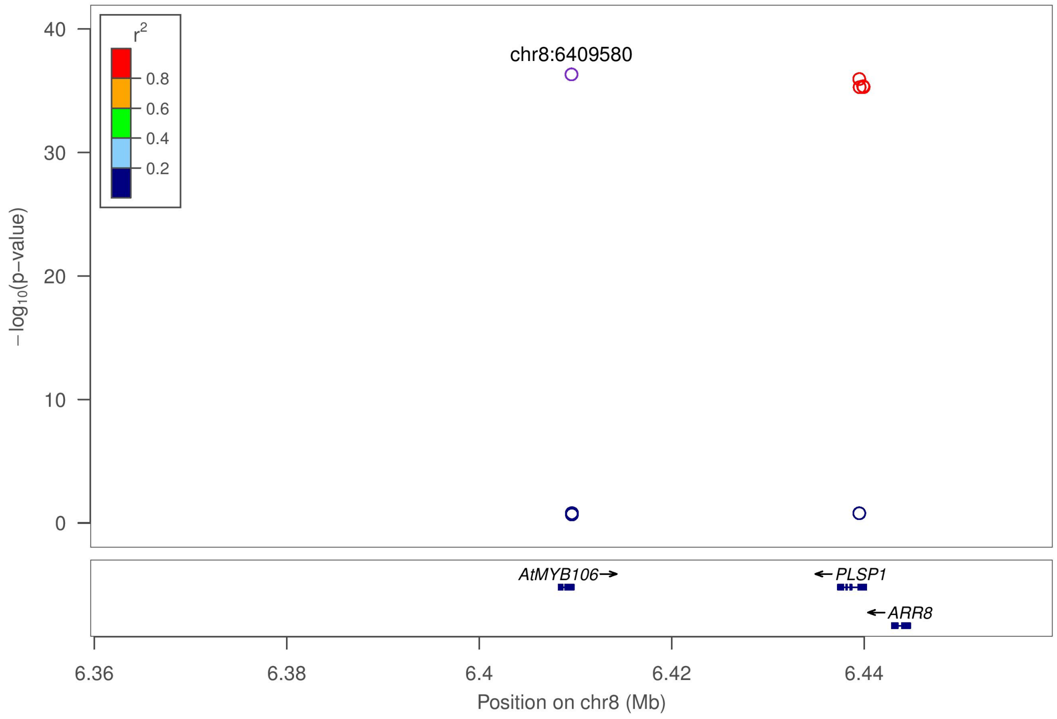


CGM severity (S12_5524524)

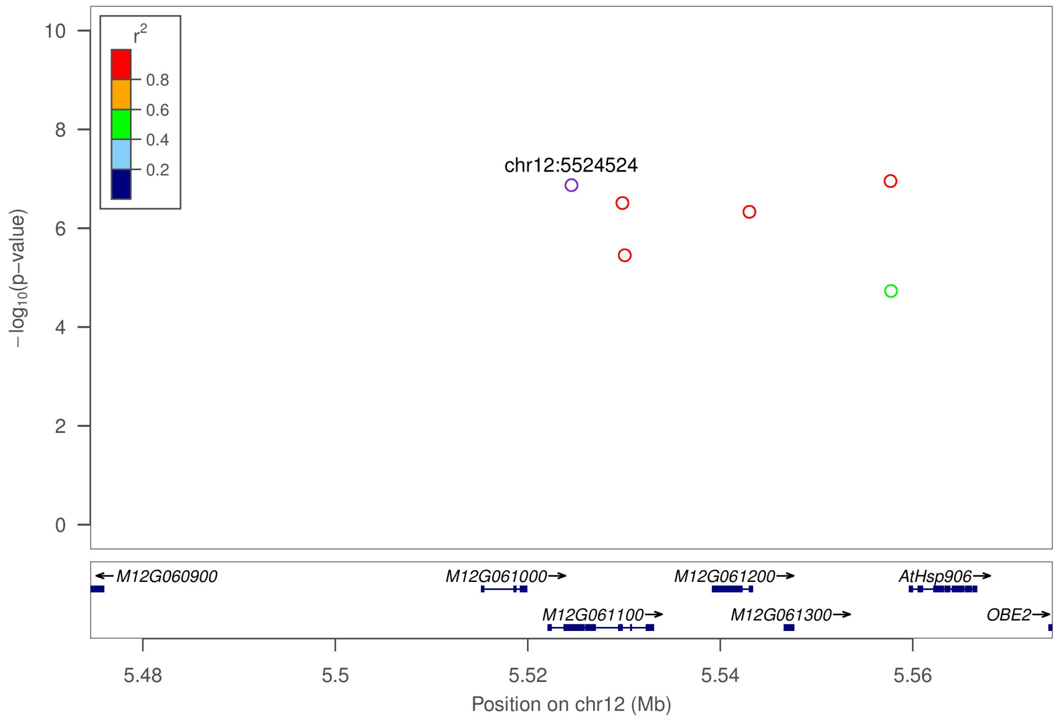

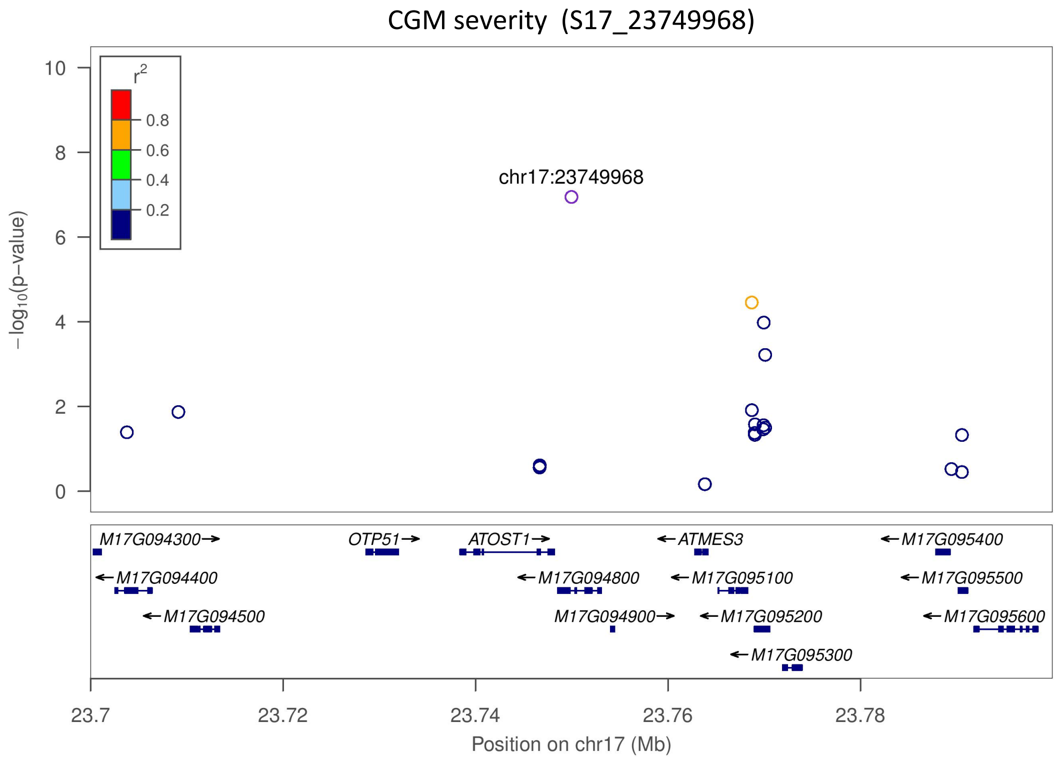

CGM severity (S17_23749968)
chr17:23749968
r2
0.8
0.6
0.4
0.2
−log10(p−value)
M17G094300
OTP51
ATOST1
ATMES3
M17G095400
M17G094400
M17G094800
M17G095100
M17G095500
M17G094500
M17G094900
M17G095200
M17G095600
M17G095300
23.7
23.72
23.74
23.76
23.78
Position on chr17 (Mb)


**C**

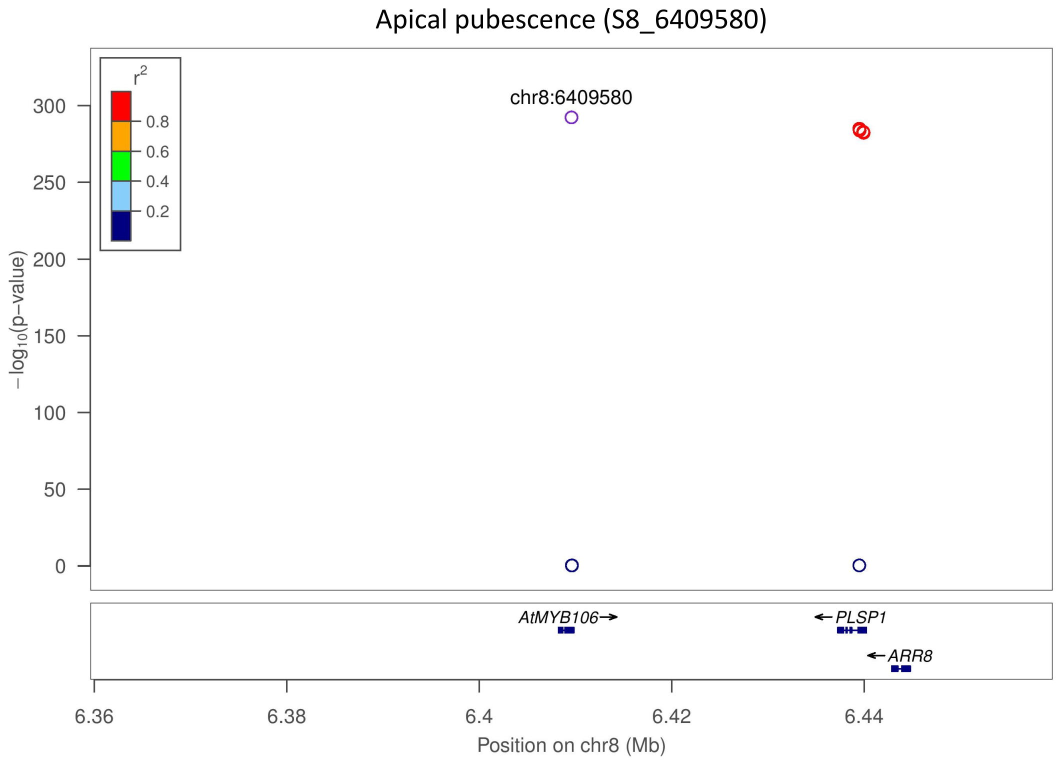

Apical pubescence (S8_6409580)
chr8:6409580
r2
0.8
0.6
0.4
0.2
−log10(p−value)
AtMYB106
PLSP1
ARR8
6.36
6.38
6.4
6.42
6.44
Position on chr8 (Mb)

### Apical pubescence (S9_1588034)

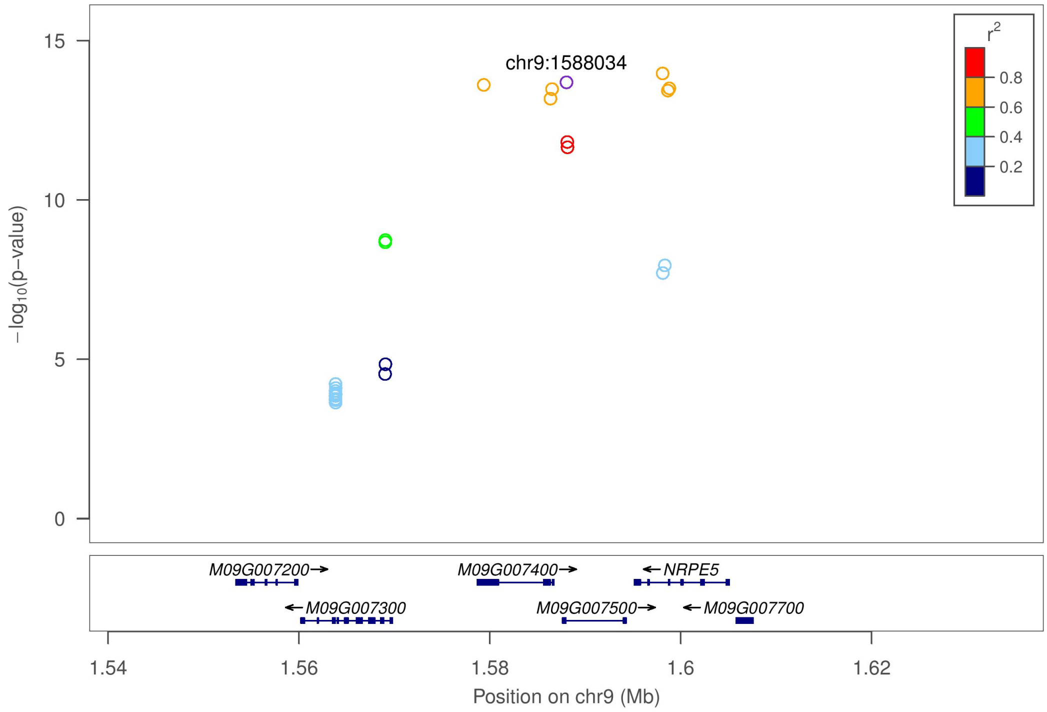


### Apical pubescence (S11_5727254)

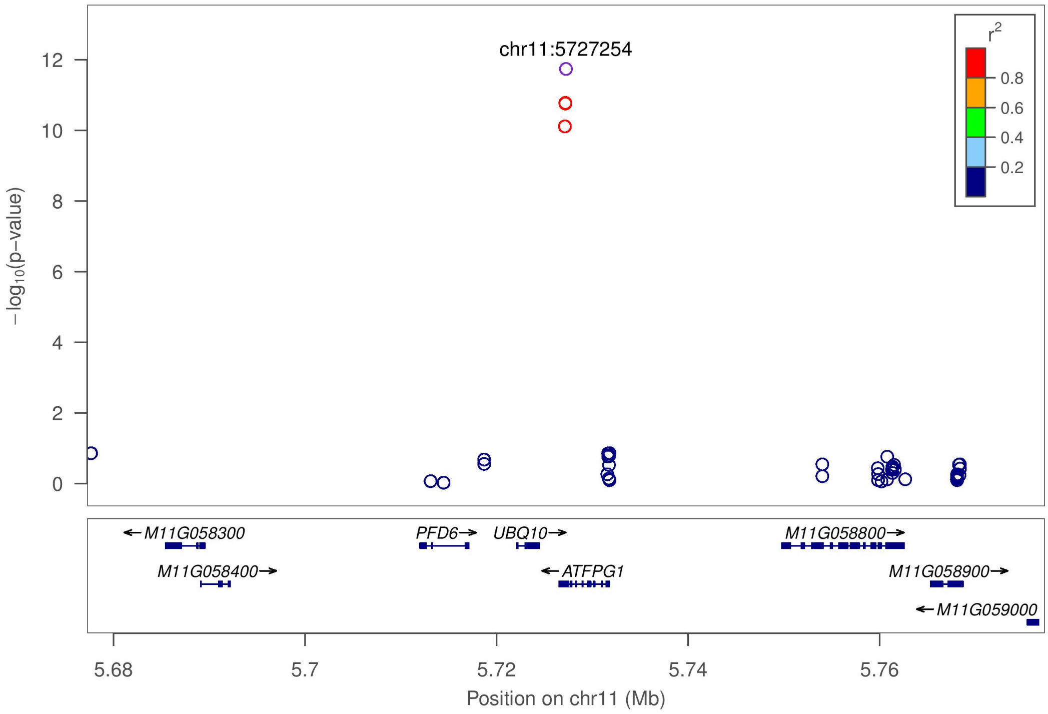

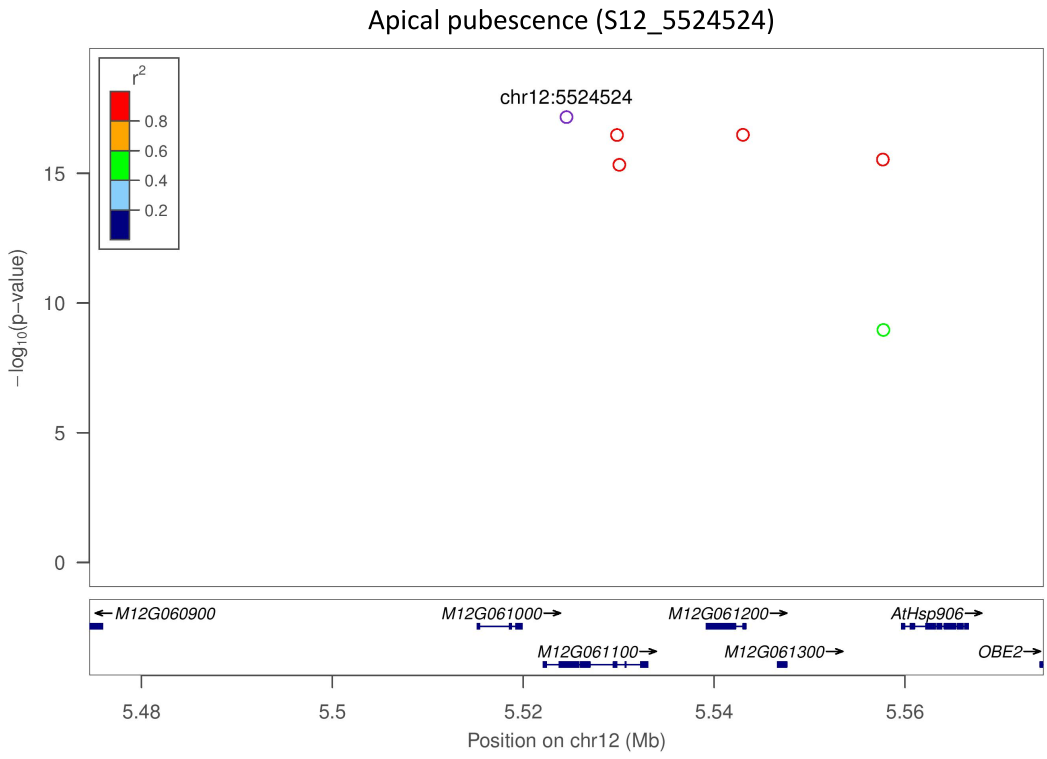

Apical pubescence (S12_5524524)
r2
0.8
0.6
0.4
0.2
chr12:5524524
15
10
5
0
−log10(p−value)
M12G060900
M12G061000
M12G061200
AtHsp906
M12G061100
M12G061300
OBE2
5.48
5.5
5.52
5.54
5.56
Position on chr12 (Mb)


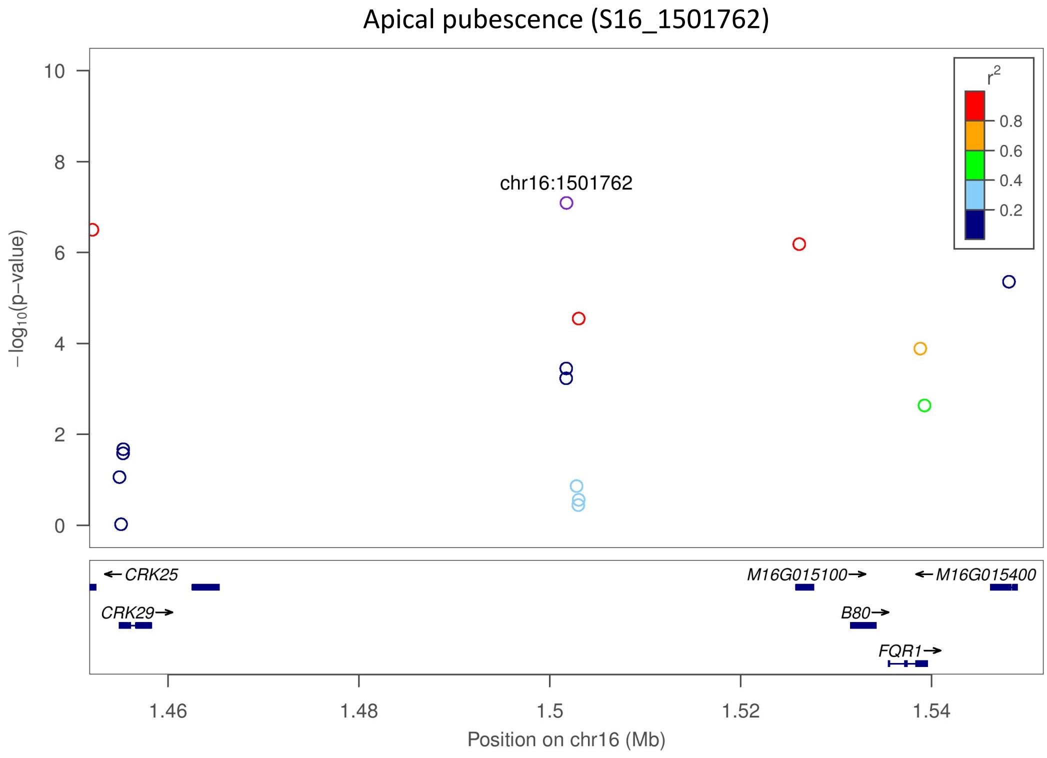

Apical pubescence (S16_1501762)
r2
0.8
0.6
0.4
0.2
chr16:1501762
10
8
6
4
2
0
−log10(p−value)
CRK25
M16G015100
M16G015400
CRK29
B80
FQR1
1.46
1.48
1.5
1.52
1.54
Position on chr16 (Mb)

**D**

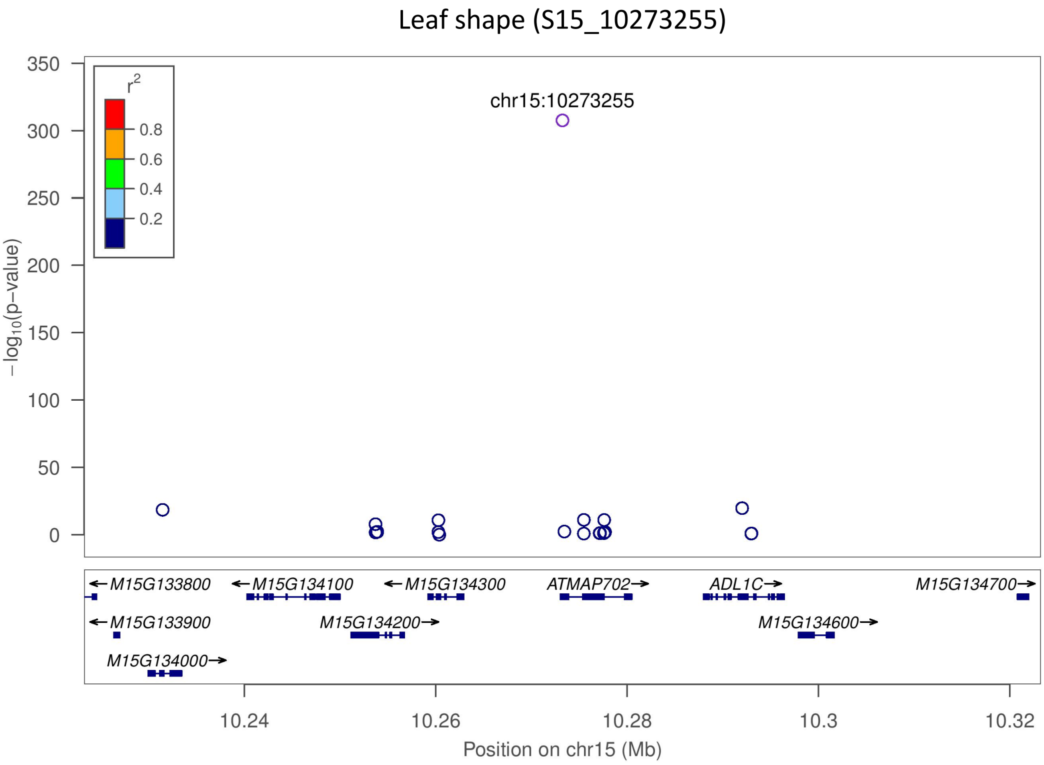
Leaf shape (S15_10273255)
chr15:10273255
r2
0.8
0.6
0.4
0.2
−log10(p−value)
350
300
250
200
150
100
50
0
M15G133800
M15G134100
M15G134300
ATMAP702
ADL1C
M15G134700
M15G133900
M15G134200
M15G134600
M15G134000
10.24
10.26
10.28
10.3
10.32
Position on chr15 (Mb)

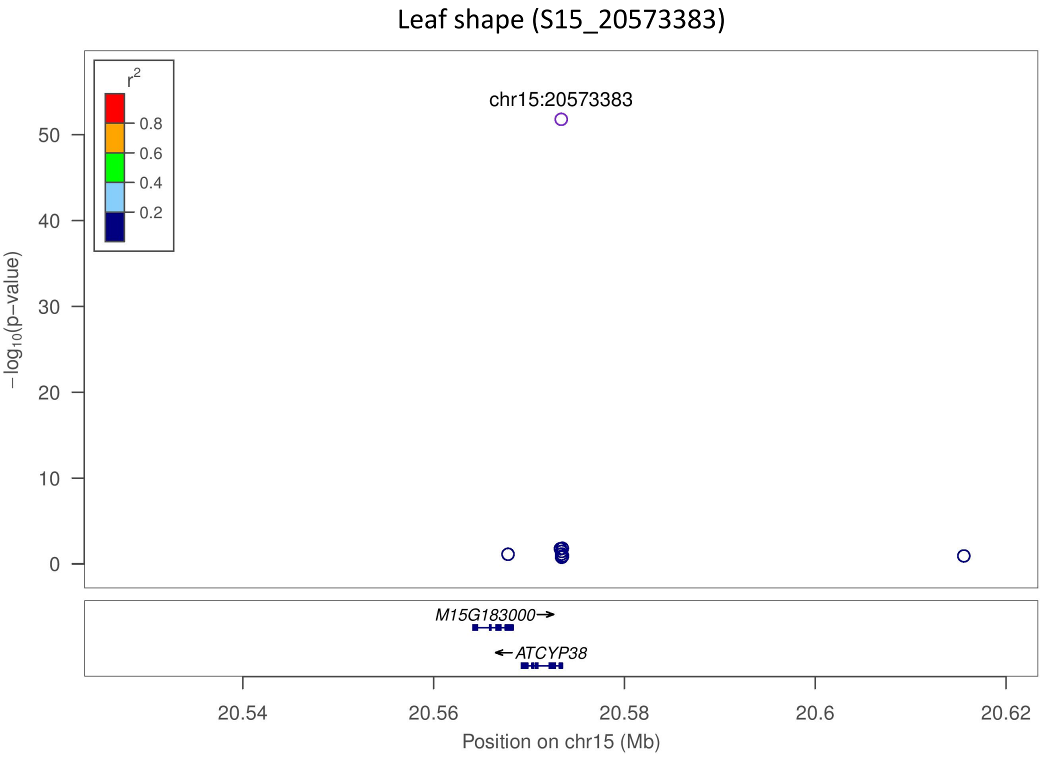
Leaf shape (S15_20573383)
chr15:20573383
r2
0.8
0.6
0.4
0.2
−log10(p−value)
50
40
30
20
10
0
M15G183000
ATCYP38
20.54
20.56
20.58
20.6
20.62
Position on chr15 (Mb)

**E**

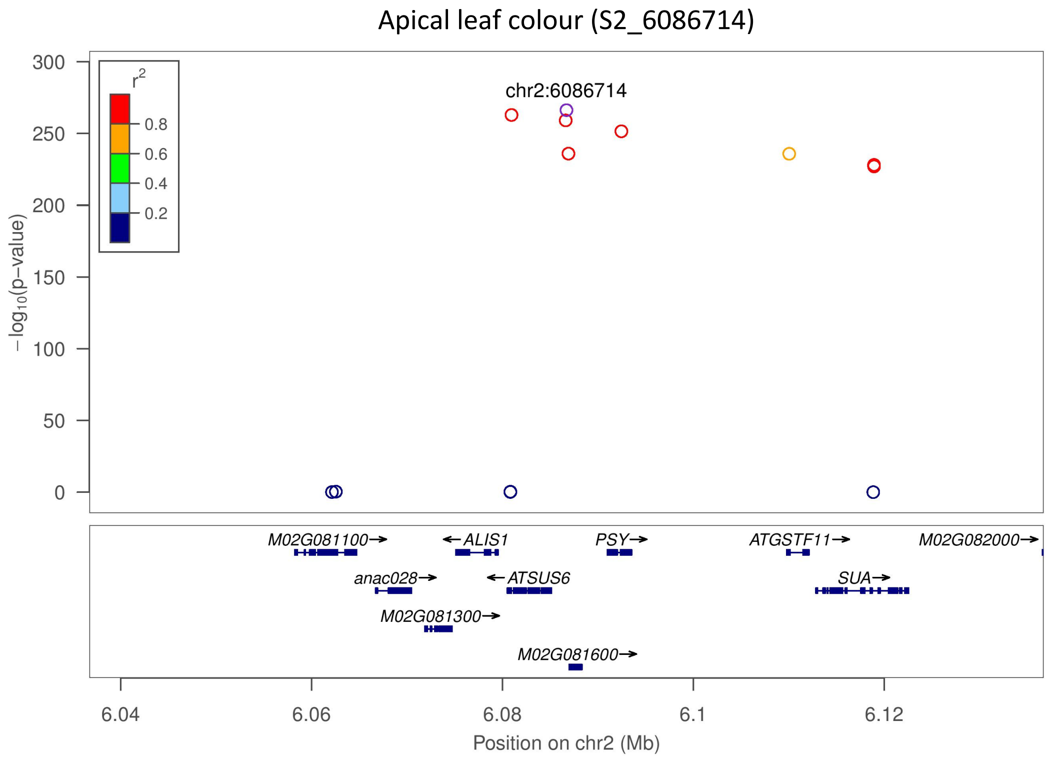

Apical leaf colour (S2_6086714)
r2
0.8
0.6
0.4
0.2
chr2:6086714
−log10(p−value)
300
250
200
150
100
50
0
M02G081100
ALIS1
PSY
ATGSTF11
M02G082000
anac028
ATSUS6
SUA
M02G081300
M02G081600
6.04
6.06
6.08
6.1
6.12
Position on chr2 (Mb)


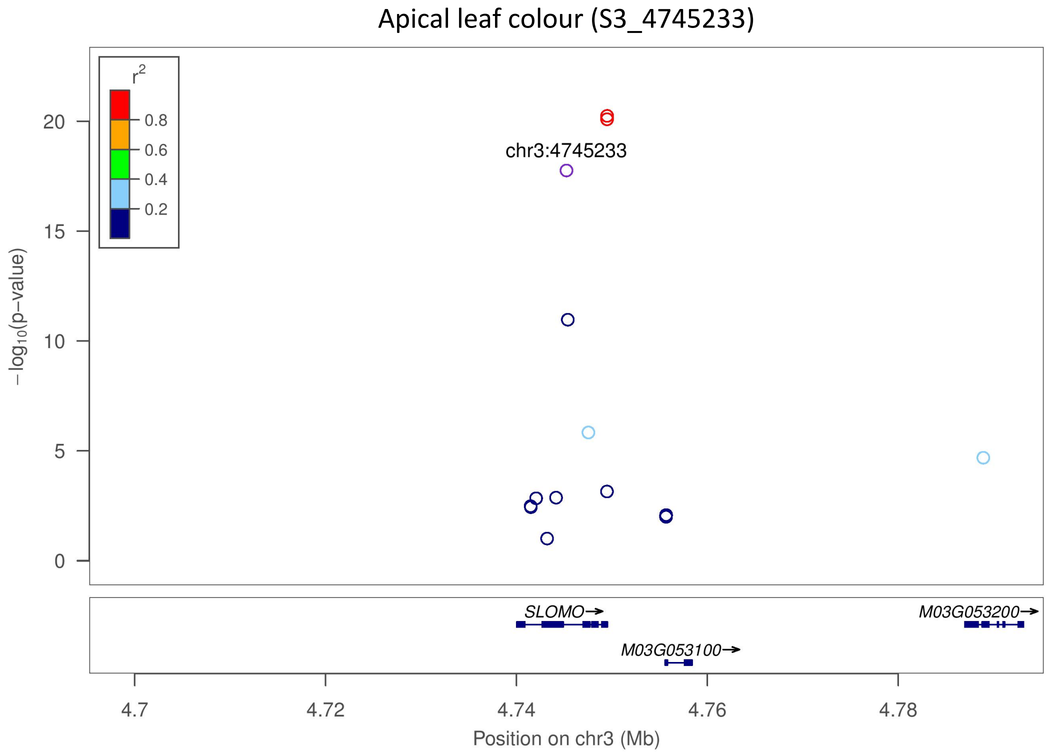

Apical leaf colour (S3_4745233)
r2
0.8
0.6
0.4
0.2
chr3:4745233
−log10(p−value)
20
15
10
5
0
SLOMO
M03G053200
M03G053100
4.7
4.72
4.74
4.76
4.78
Position on chr3 (Mb)

Apical leaf colour (S8_6061421)

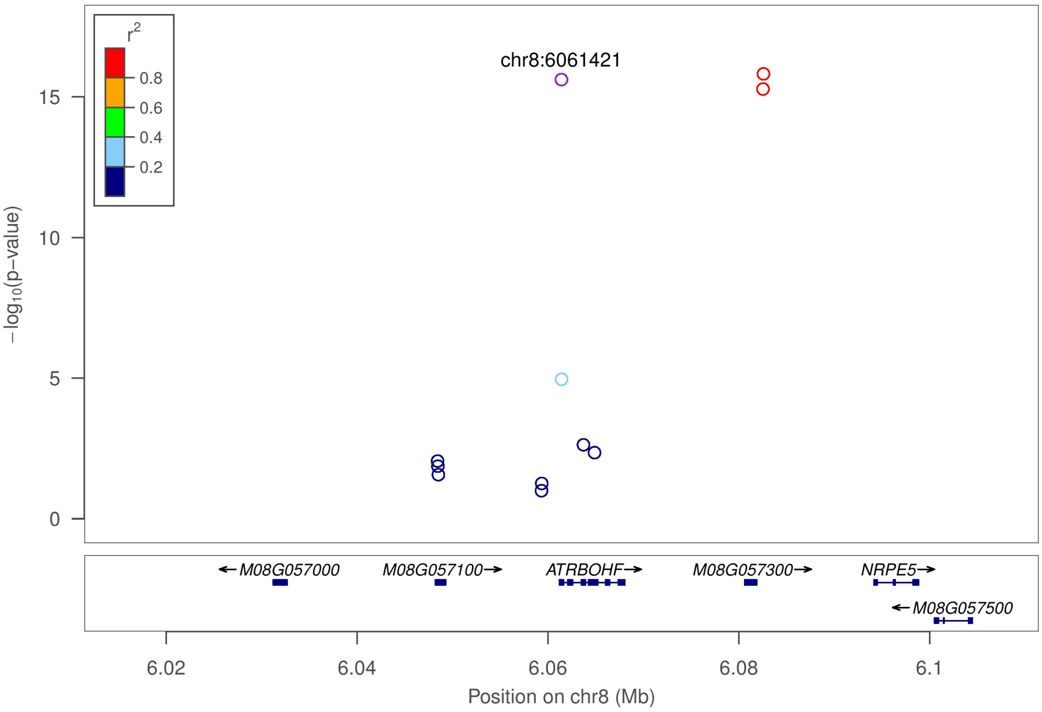


**F**

Mature leaf greenness (S1_23452638)

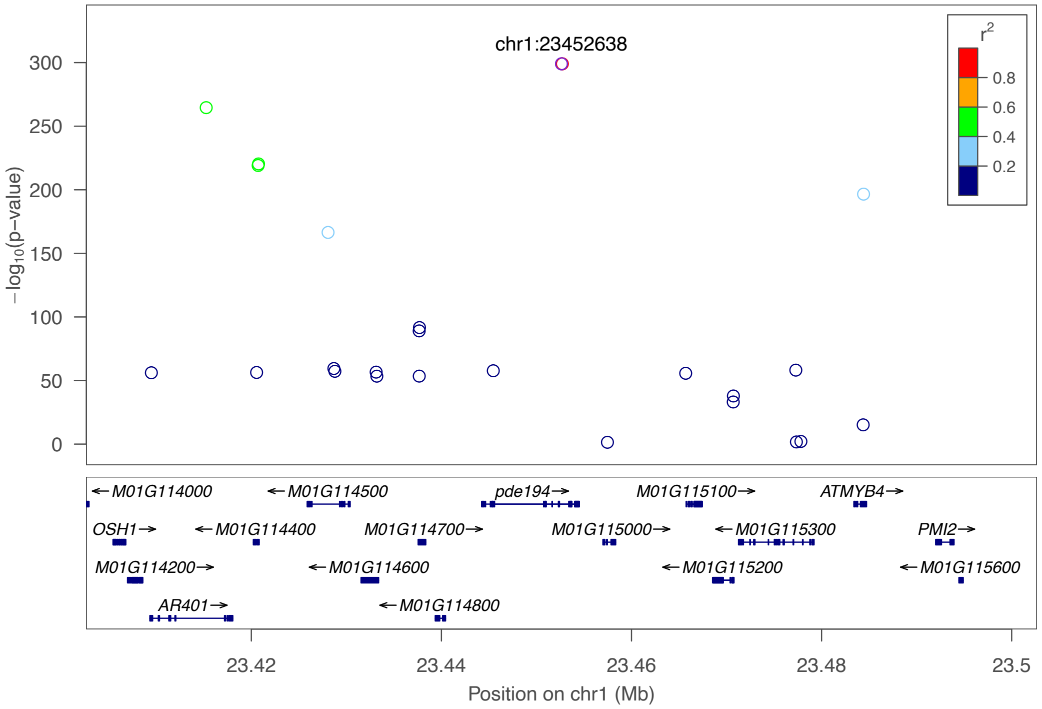

**G**

Petiole colour (S1_23055068)

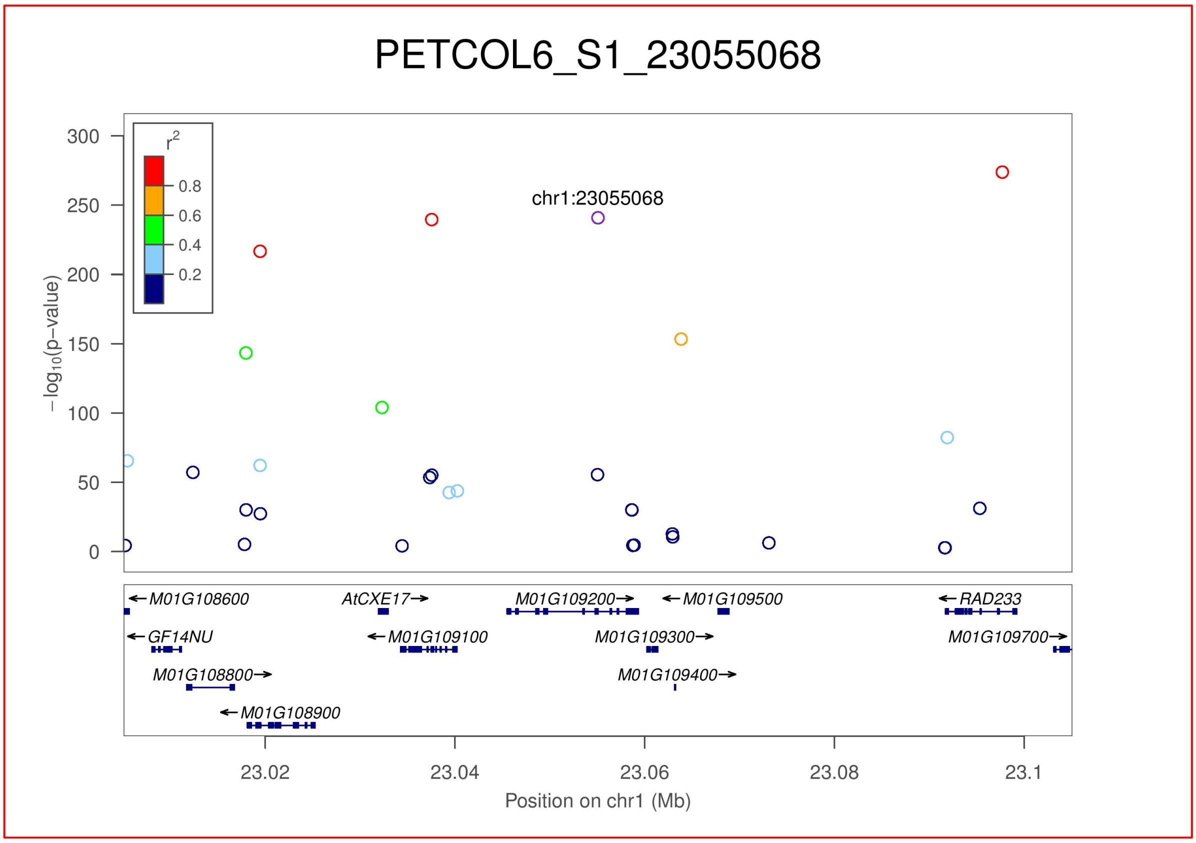


**H**

Harvest index (S2_2809137)

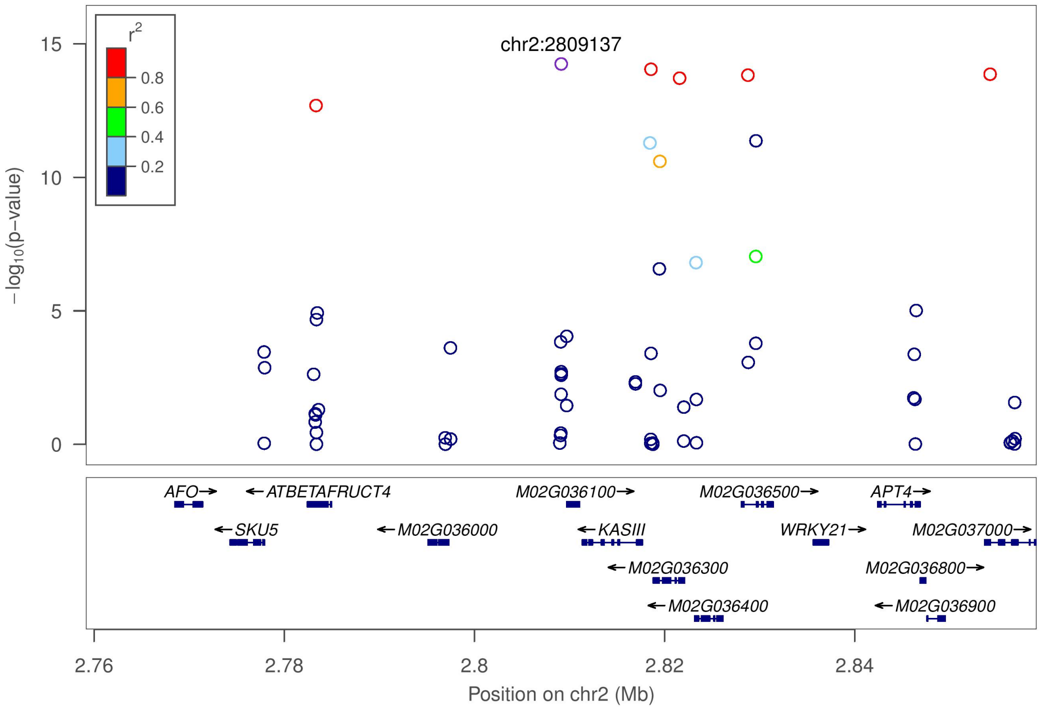

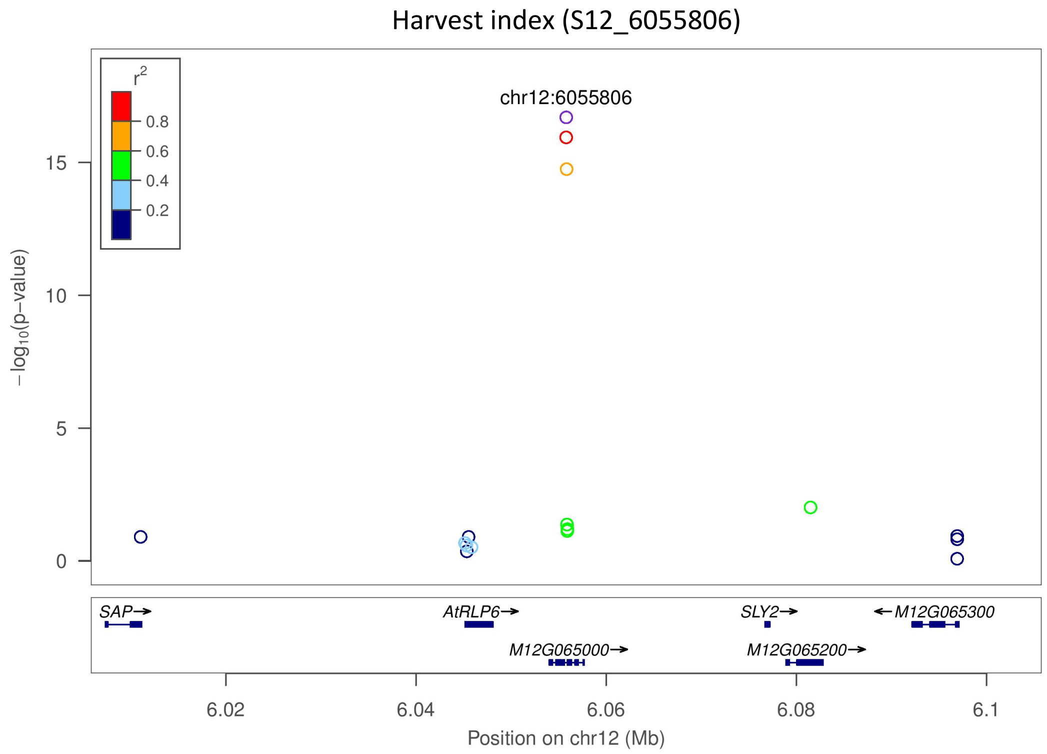
Harvest index (S12_6055806)
r2
0.8
0.6
0.4
0.2
chr12:6055806
−log10(p−value)
15
10
5
0
SAP
AtRLP6
M12G065000
SLY2
M12G065200
M12G065300
6.02
6.04
6.06
6.08
6.1
Position on chr12 (Mb)

I

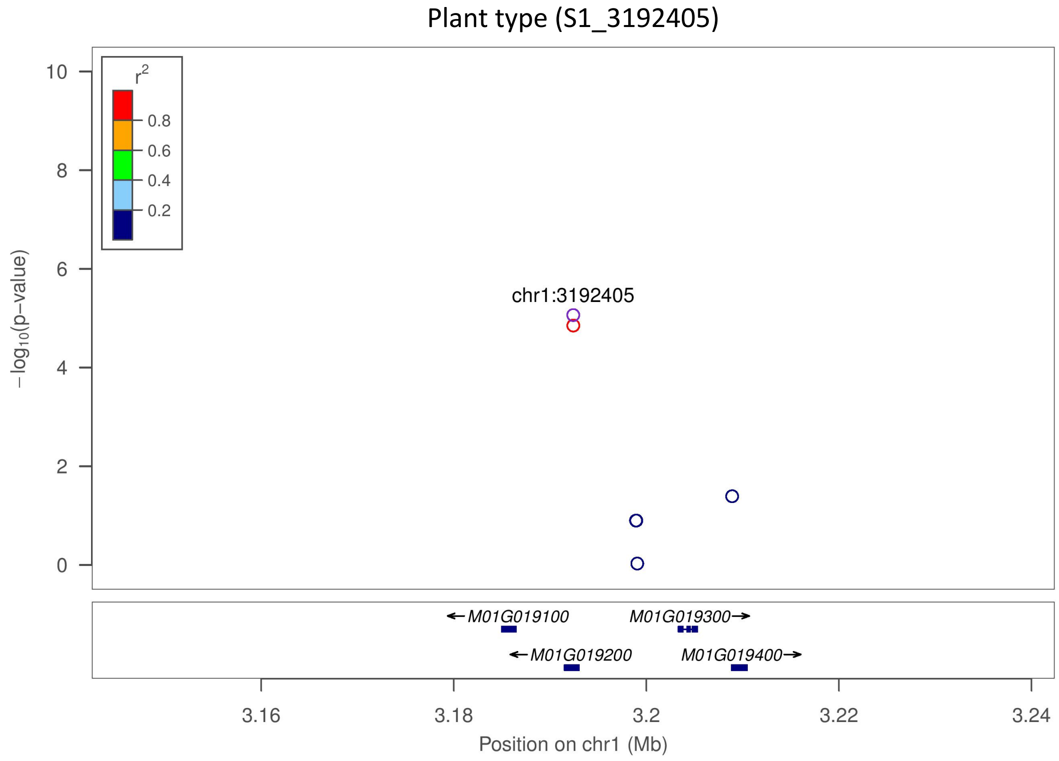
Plant type (S1_3192405)
r2
0.8
0.6
0.4
0.2
chr1:3192405
−log10(p−value)
10
8
6
4
2
0
M01G019100
M01G019300
M01G019200
M01G019400
3.16
3.18
3.2
3.22
3.24
Position on chr1 (Mb)

Plant type (S1_25303195)

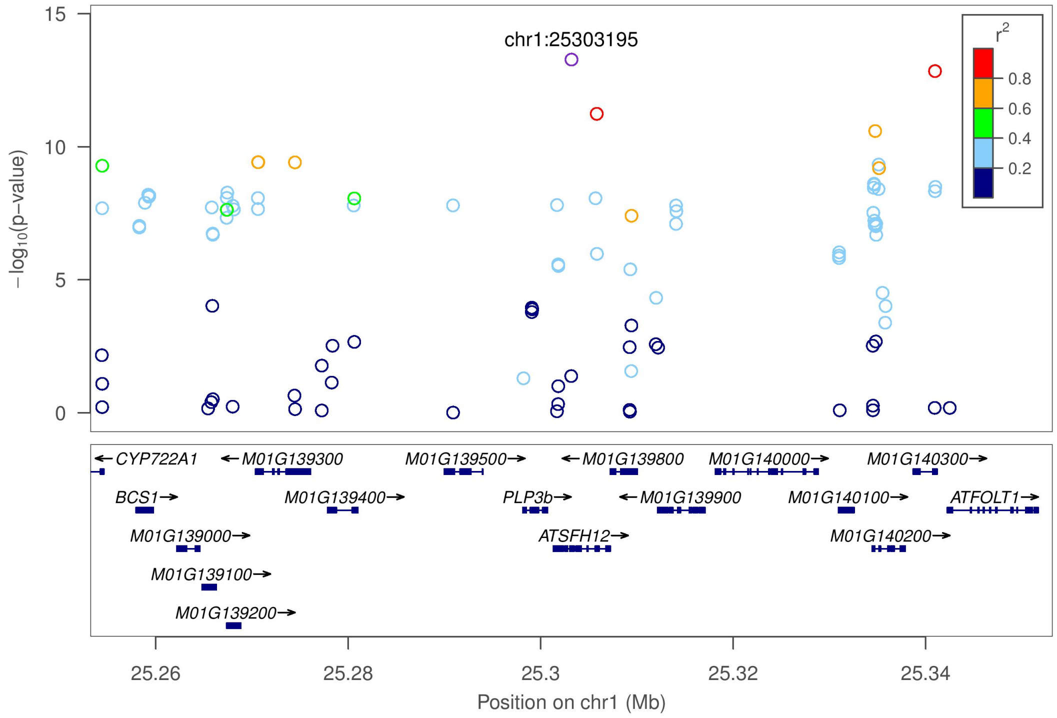


J

Outer stem colour (S2_13928566)

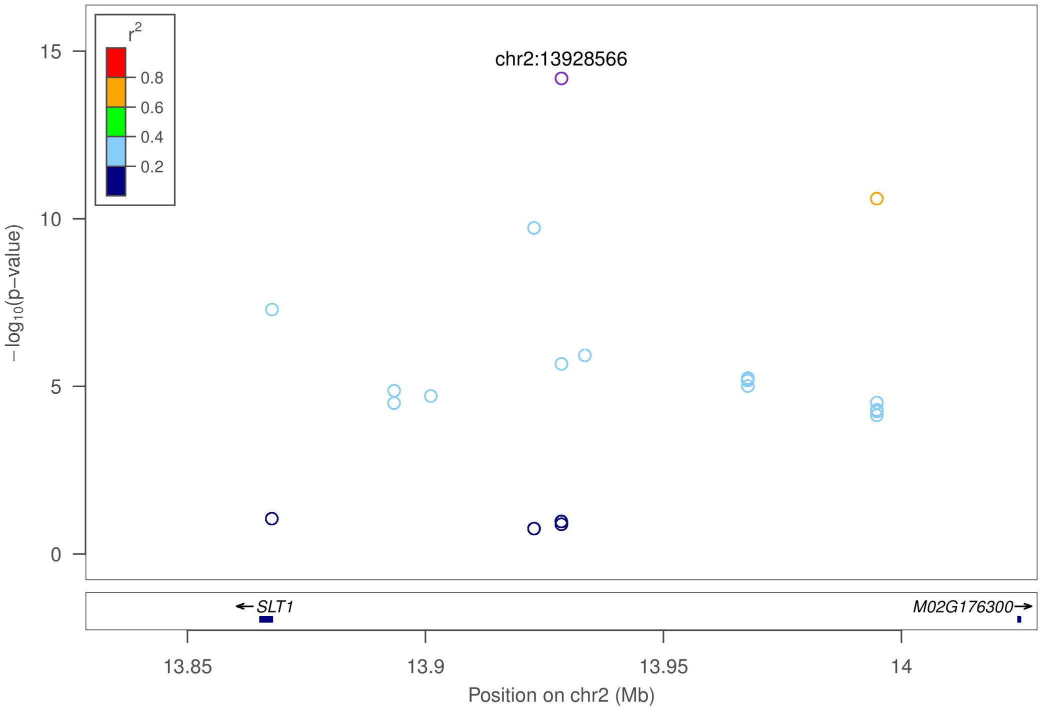

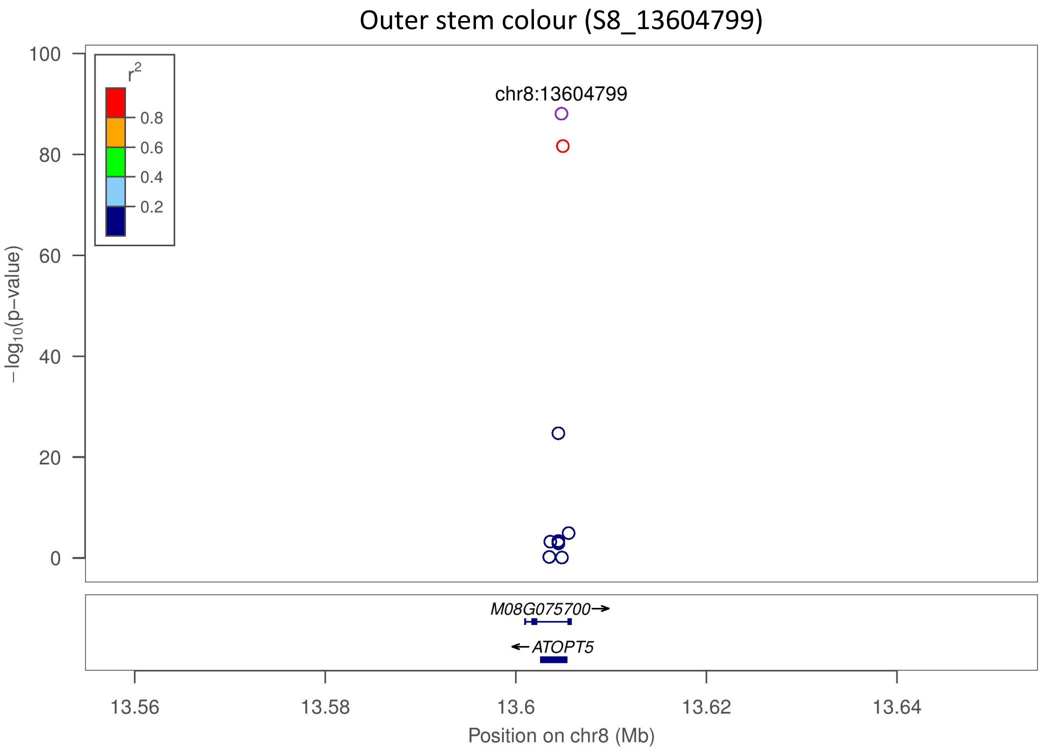
Outer stem colour (S8_13604799)
chr8:13604799
r2
0.8
0.6
0.4
0.2
−log10(p−value)
M08G075700→
←ATOPT5
13.56
13.58
13.6
13.62
13.64
Position on chr8 (Mb)

**K**

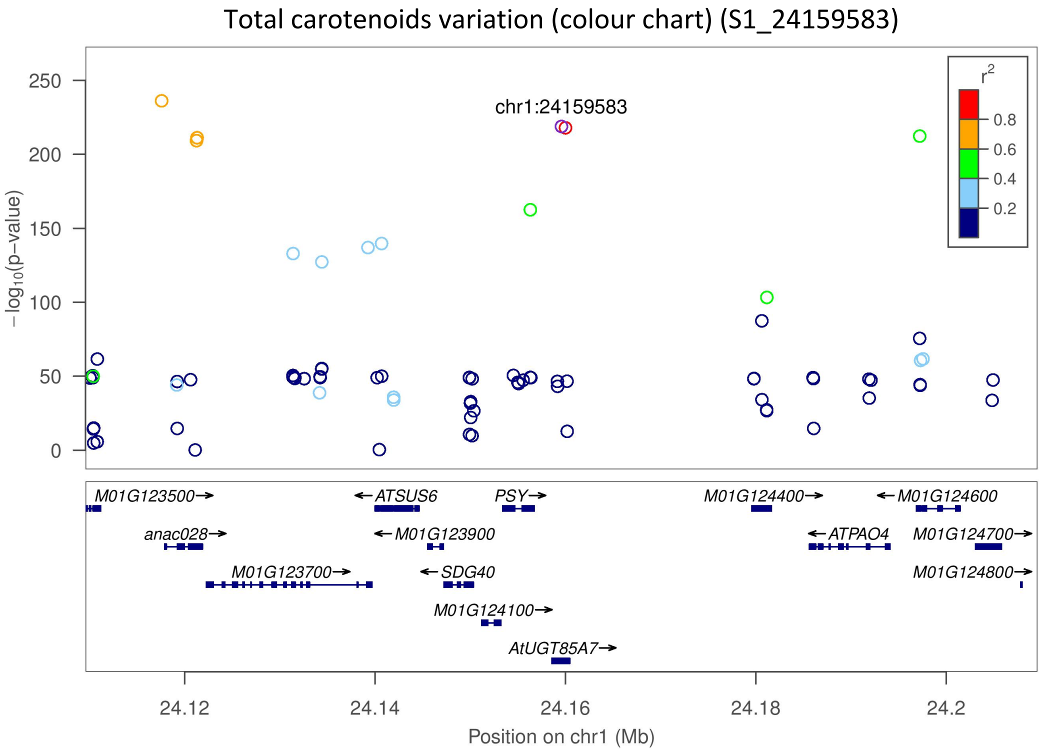
Total carotenoids variation (colour chart) (S1_24159583)
chr1:24159583
r2
0.8
0.6
0.4
0.2
−log10(p−value)
M01G123500→
←ATSUS6
PSY→
M01G124400→
←M01G124600
anac028→
←M01G123900
←ATPAO4
M01G124700→
M01G123700→
←SDG40
M01G124800→
M01G124100→
AtUGT85A7→
24.12
24.14
24.16
24.18
24.2
Position on chr1 (Mb)

Total carotenoids variation (colour chart) (S1_24636113)

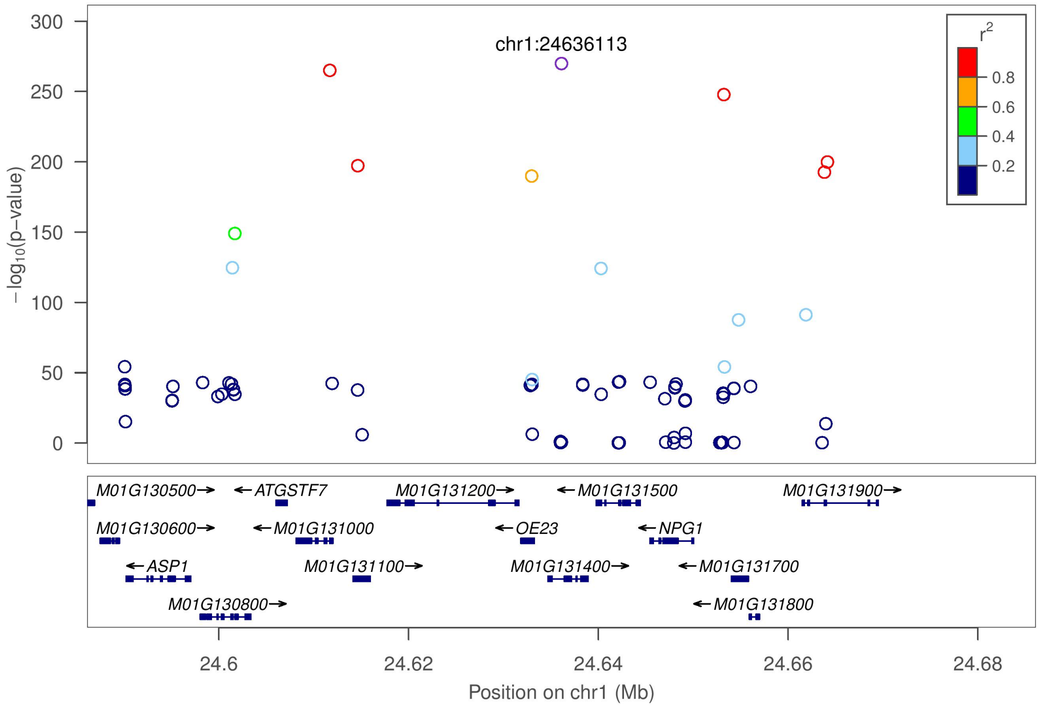


Total carotenoids variation (colour chart) (S1_30543962)

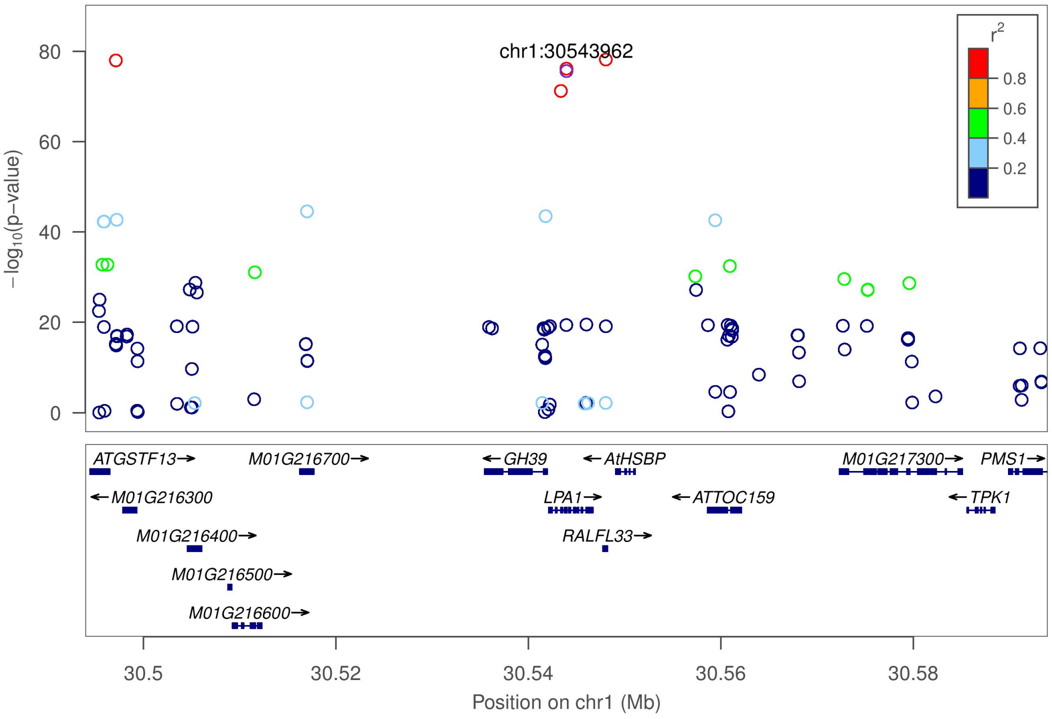

Total carotenoids variation (colour chart) (S5_3387558)

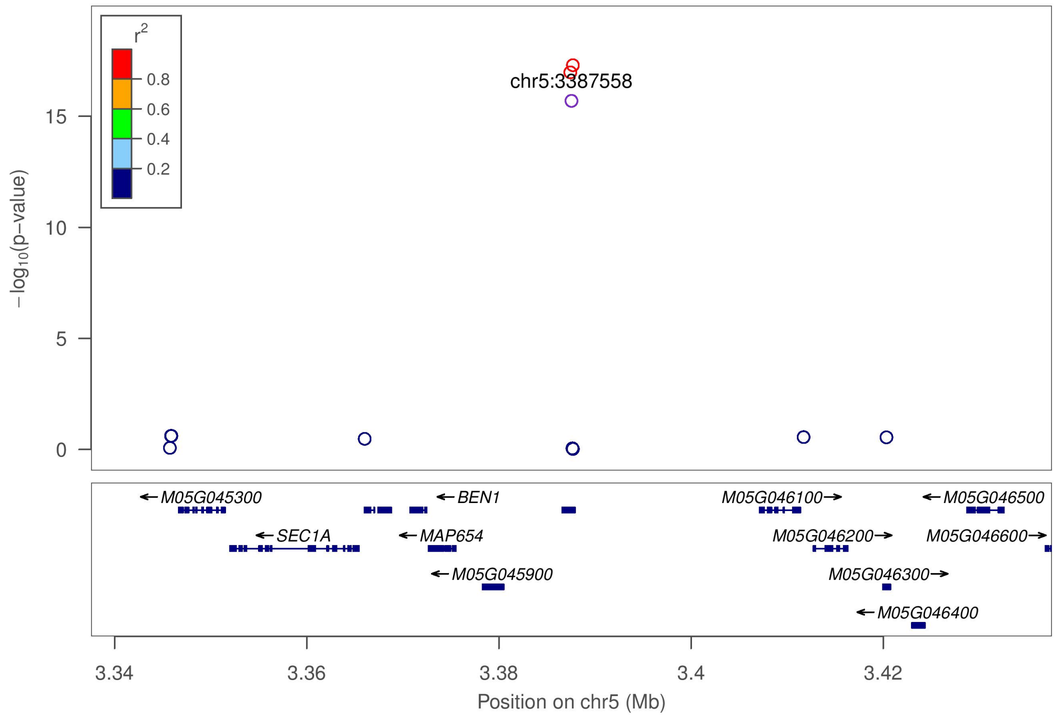


Total carotenoids (colour chart) (S8_4319215)

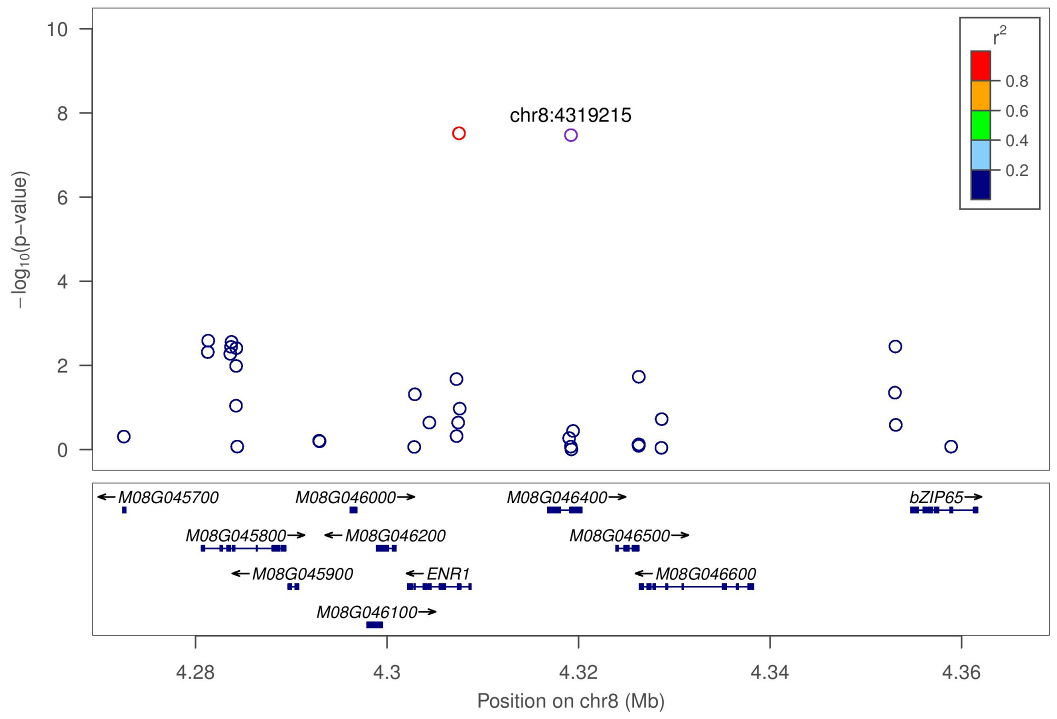

Total carotenoids variation (colour chart) (S8_25598183)

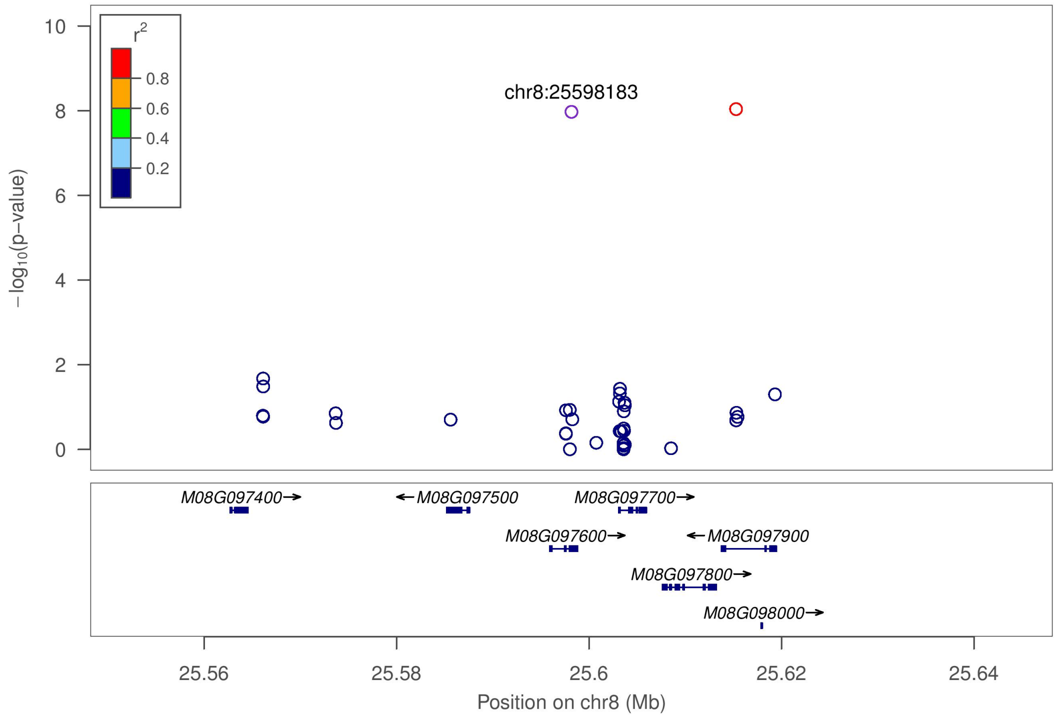


Total carotenoids variation (colour chart) (S15_7659426)

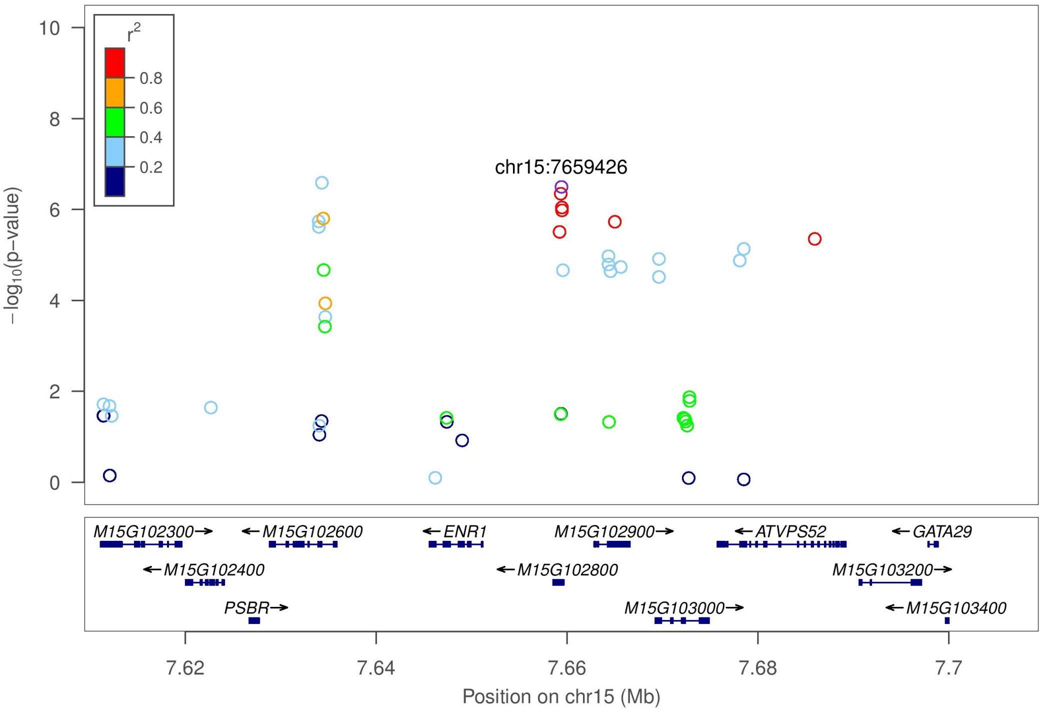

Total carotenoids variation (colour chart) (S16_484011)

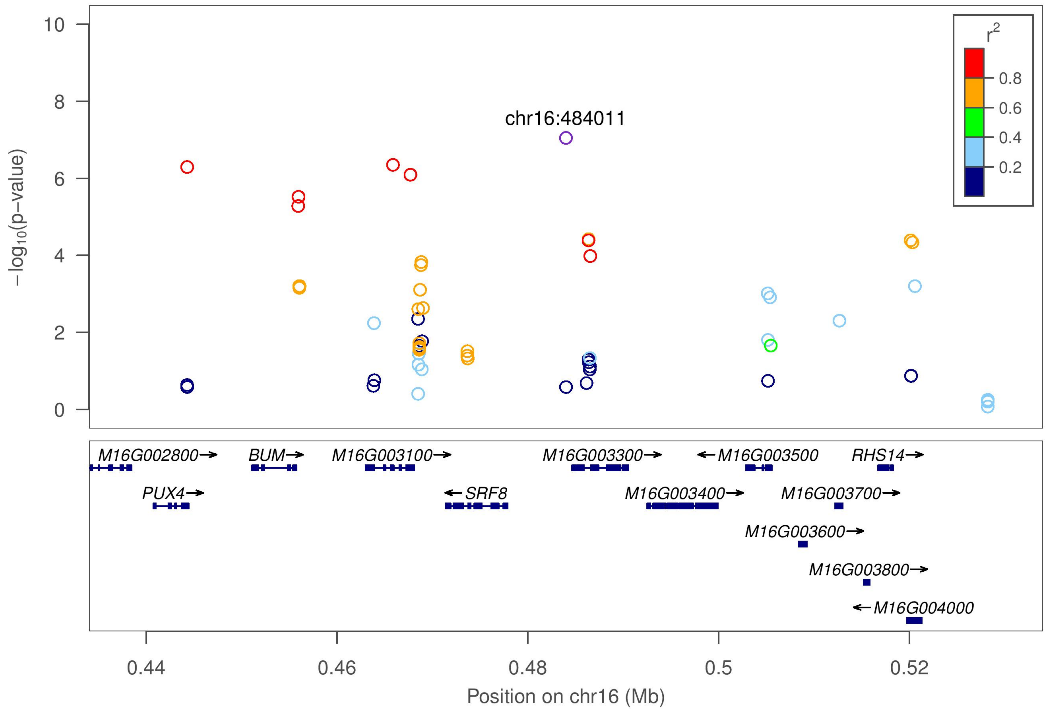


**L**

Dry matter content (S1_24636113)

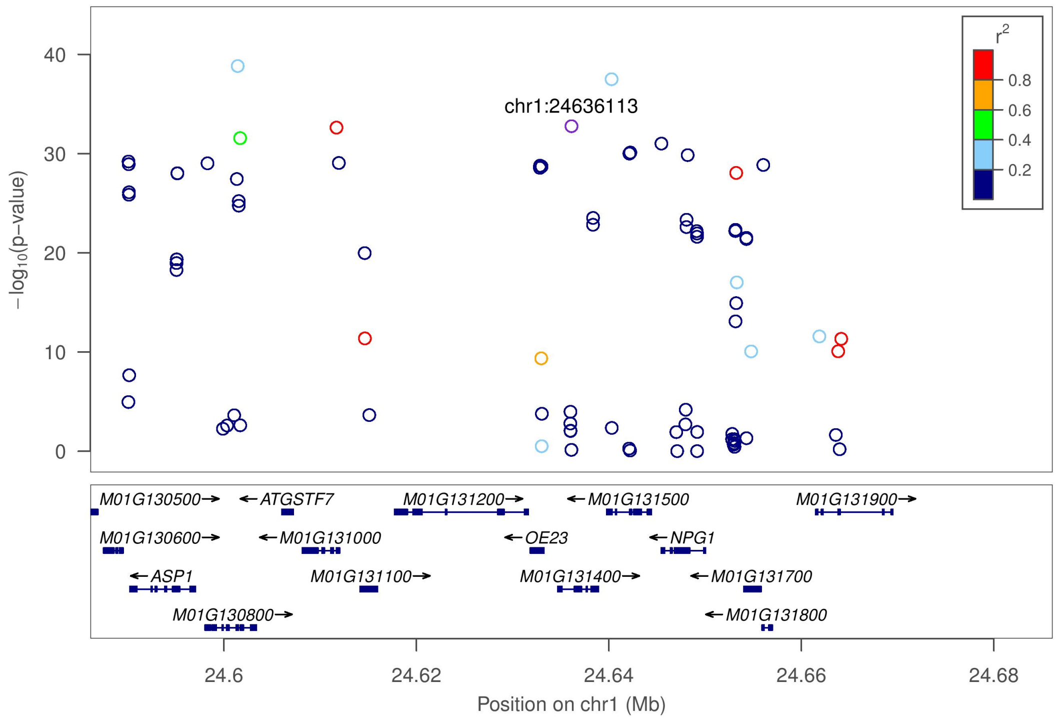

Dry matter content (S6_20589894)

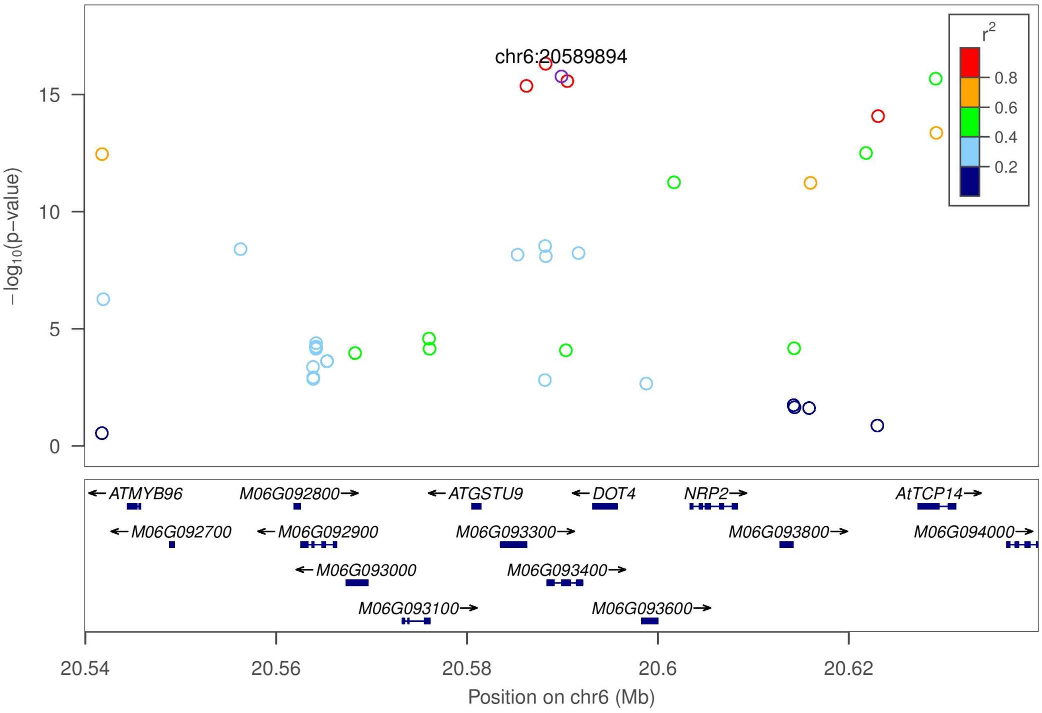


Dry matter content (S12_5524524)

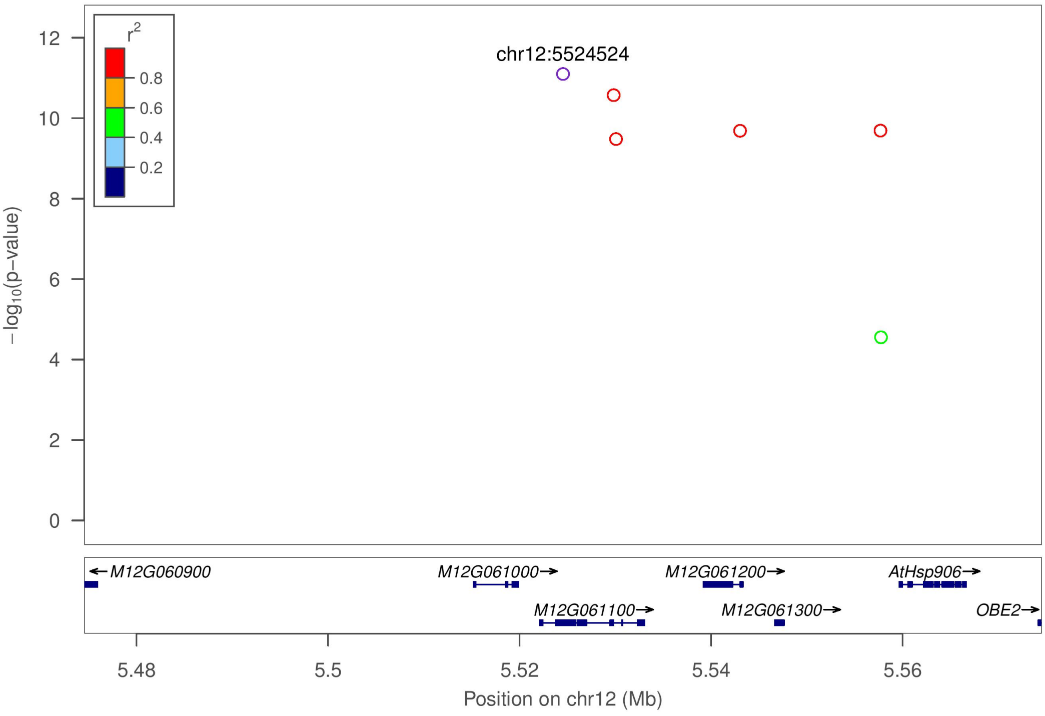

Dry matter content (S15_1012346)

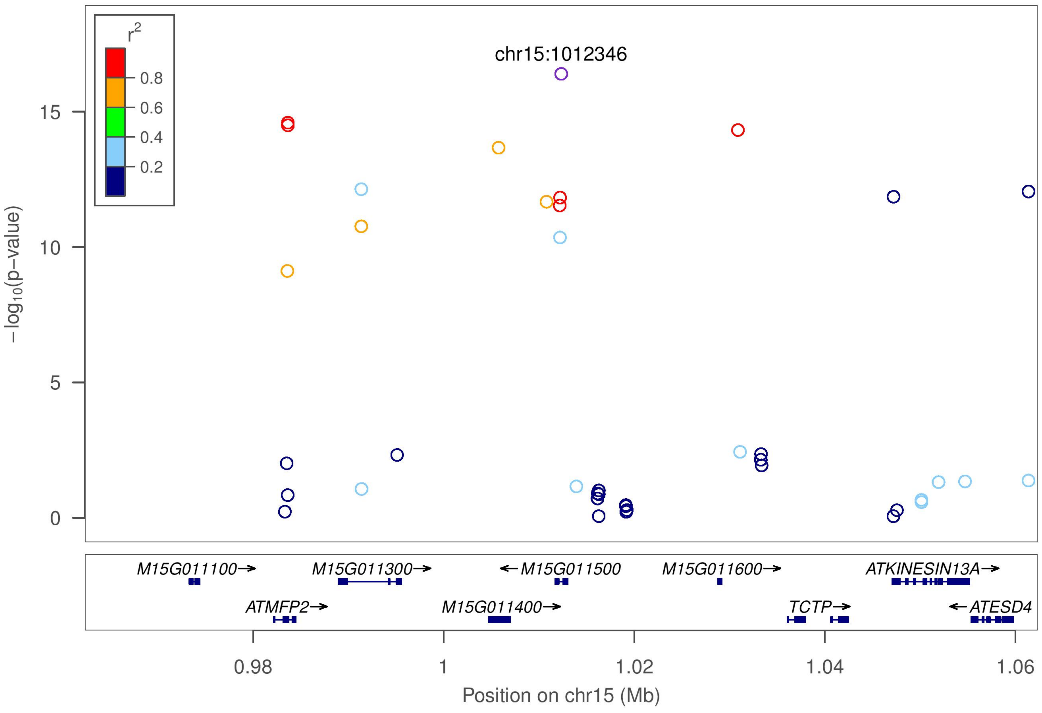


Dry matter content (S16_25663808)

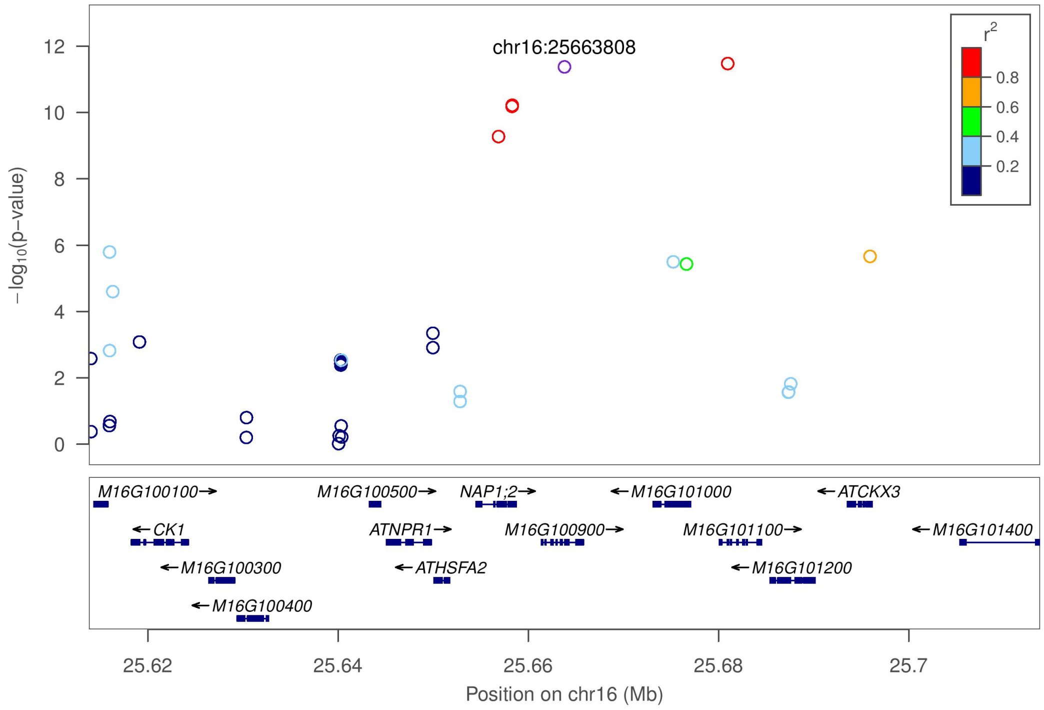

**M**

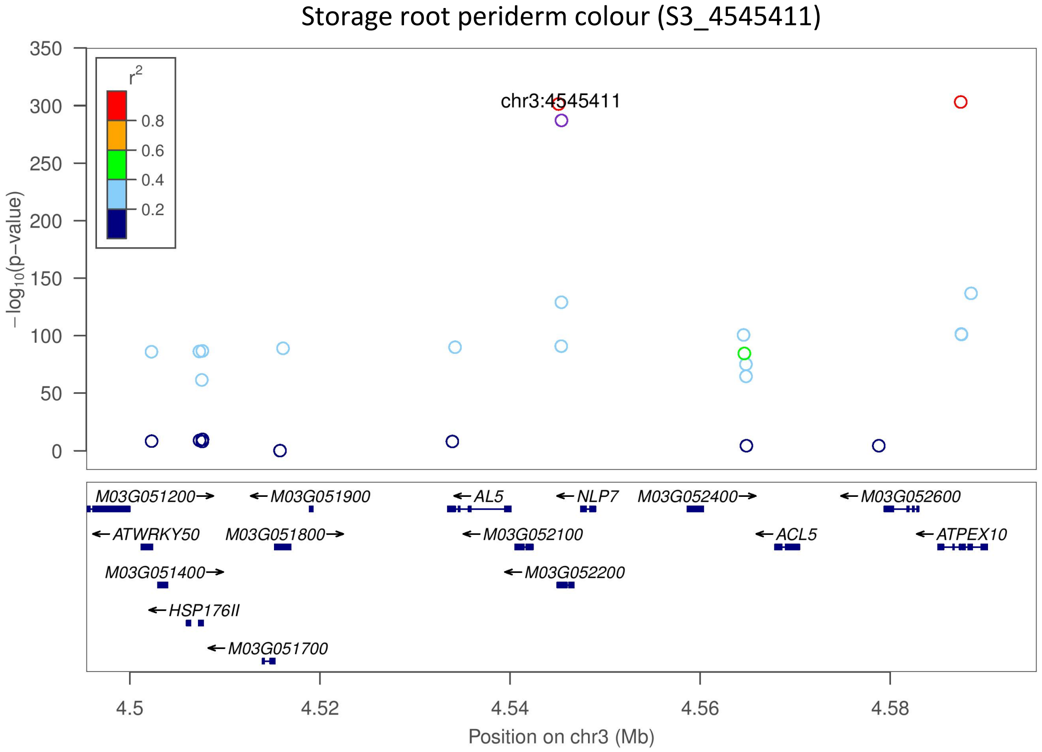

Storage root periderm colour (S3_4545411)
r2
0.8
0.6
0.4
0.2
chr3:4545411
−log10(p−value)
M03G051200
M03G051900
AL5
NLP7
M03G052400
M03G052600
ATWRKY50
M03G051800
M03G052100
ACL5
ATPEX10
M03G051400
M03G052200
HSP176II
M03G051700
Position on chr3 (Mb)


**N**

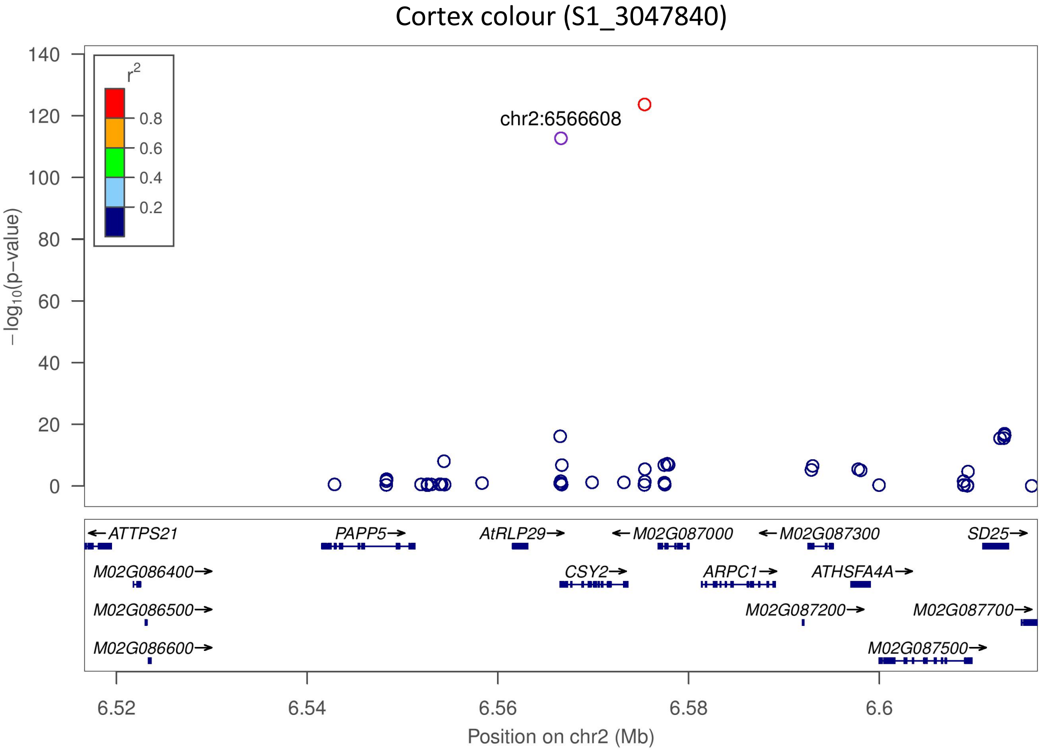

Cortex colour (S1_3047840)
r2
0.8
0.6
0.4
0.2
chr2:6566608
−log10(p−value)
ATTPS21
PAPP5
AtRLP29
M02G087000
M02G087300
SD25
M02G086400
CSY2
ARPC1
ATHSFA4A
M02G086500
M02G087200
M02G087700
M02G086600
M02G087500
Position on chr2 (Mb)

Cortex colour (S2_6566608)

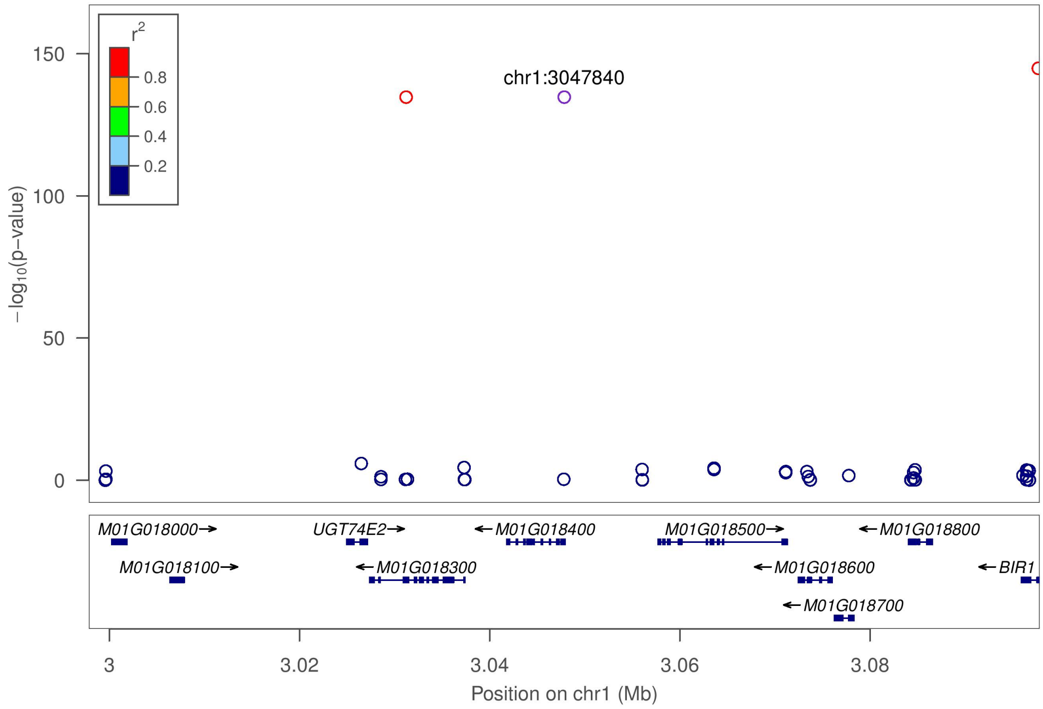


**Figure S4** Regional Manhattan plots for each locus-trait combination. Plots include candidate genes within 100 Kb of the top SNP marker (50 kb upstream, 50 kb downstream) with some adjustments based on the extent of local linkage disequilibrium with the candidate marker. SNPs are color coded based on linkage disequilibrium with the top marker. (A) CMD severity; (B) CGM severity; (C) Apical leaf pubescence; (D) Leaf shape; (E) Apical leaf colour; (F) Mature leaf greenness; (G) Petiole colour; (H) Harvest index; (I) Plant type; (J) Outer stem colour; (K) Total carotenoids content; (L) Dry matter content; (M) Periderm colour; (N) Root cortex colour.
